# Visualizing structural transitions of ligand-dependent gating of the TRPM2 channel

**DOI:** 10.1101/516468

**Authors:** Ying Yin, Mengyu Wu, Allen L. Hsu, William F. Borschel, Mario J. Borgnia, Gabriel C. Lander, Seok-Yong Lee

**Affiliations:** Department of Biochemistry, Duke University School of Medicine, Durham, North Carolina, 27710, USA.; Department of Integrative Structural and Computational Biology, The Scripps Research Institute, La Jolla, California, 92037, USA.; Genome Integrity and Structural Biology Laboratory, National Institute of Environmental Health Sciences, National Institutes of Health, Department of Health and Human Services, Research Triangle Park, NC 27709, USA.

## Abstract

The calcium-permeable transient receptor potential melastatin 2 (TRPM2) channel plays a key role in redox sensation in many cell types ^1–3^. Channel activation requires binding of both ADP-ribose (ADPR) ^2,4–6^ and Ca^2+ 7^. The recently published TRPM2 structures from *Danio rerio* in the ligand-free and in the ADPR/Ca^2+^-bound conditions represent the channel in closed and open states, which uncover substantial tertiary and quaternary conformational rearrangements ^8^. However, it is unclear how these rearrangements occur within the tetrameric channel during channel gating. Here we report two cryo-electron microscopy structures of TRPM2 from the same species in complex with Ca^2+^ alone, and with both ADPR and Ca^2+^, determined to an overall resolution of ~3.8 Å and ~4.2 Å respectively. In comparison with the published results, our studies capture TRPM2 in two-fold symmetric intermediate states, offering a glimpse of the structural transitions within the tetramer that bridge the closed and open conformations.

---

The transient receptor potential melastatin (TRPM) ion channel family is part of the TRP channel superfamily and comprises eight members (TRPM1 to TRPM8) that carry out diverse functions in a variety of physiological pathways ^9^. TRPM2 is a calcium-permeable non-selective cation channel that is widely expressed in the nervous, immune, and endocrine systems, and plays a crucial role in warmth and redox-dependent signaling ^1–3^. Studies of TRPM2-deficient mice have shown that TRPM2 channels expressed in sensory and central neurons are responsible for sensation of warm temperatures and body temperature regulation ^10,11^. TRPM2 has also been found to be activated by reactive oxygen species (ROS), consequently playing a key role in Ca^2+^ signaling involved in chemokine production, insulin production, and cell death under oxidative stress ^12–16^.

Extensive electrophysiological studies have shown that activation of TRPM2 by ROS is caused by an increase of intracellular ADP-ribose (ADPR), a TRPM2 agonist, and that both ADPR and Ca^2+^ are required for channel activation ^2,4–7,17^. Notably, TRPM2 contains a putative enzyme domain, called the Nudix Hydrolase 9 Homology (NUDT9H) domain located at the C-terminus, which exhibits a high degree of homology to the mitochondrial ADP-ribose pyrophosphatase NUDT9 ^18,19^. As such, TRPM2 was initially classified as a channel-enzyme in which enzymatic activity was believed to be coupled to channel gating ^2,18^. However, subsequent studies have since repudiated this mechanism, as the NUDT9H domain lacks enzymatic activity. Instead it has been suggested that the NUDT9H domain merely serves as a binding site for ADPR ^20^.

Recently, structures of the zebrafish *Danio rerio* TRPM2 were reported in the ligand-free closed state and in the ADPR/Ca^2+^-bound open state (hereafter referred to as TRPM2_DR_closed_ and TRPM2_DR_open_ respectively) ^8^. This study not only revealed the location of the NUDT9H domain, but also showed that the Ca^2+^ ion binds in the cavity formed by the voltage-sensor like domain (VSLD) comprising transmembrane segments 1-4 (S1-S4). Unexpectedly, ADPR was found to bind in the cleft of the melastatin homology domain 1/2 (MHR1/2) and not, as had long been predicted, in the NUDT9H domain. Structural analyses of the closed and open conformations have provided a mechanism for ligand-dependent activation of the channel, wherein binding of ADPR in the MHR1/2 domain triggers substantial conformational rearrangements in the cytoplasmic domain (CD), which are sequentially transduced and propagated to the distal transmembrane channel domain (TMD) to induce pore opening.

In spite of this progress towards understanding ligand-dependent activation of the TRPM2 channel, it remains unclear whether these drastic quaternary structural rearrangements occur in a concerted, four-fold symmetric manner. We recently reported that the transient receptor potential vanilloid 2 (TRPV2) channel adopts two-fold symmetric conformations upon resiniferatoxin (RTx)-mediated activation ^21^, and that two-fold symmetric states are associated with ligand-dependent gating of the TRPV3 channel ^22^. It is presently unclear if reduced symmetry is associated with gating in other TRP channel families. To address these questions, we determined the cryo-electron microscopy (cryo-EM) structures of the full-length TRPM2 from *Danio rerio* (TRPM2_DR_) in the presence of Ca^2+^ (referred to as TRPM2_DR_Ca2+_) and in the presence of both ADPR and Ca^2+^ (referred to as TRPM2_DR_ADPR/Ca2+_). Our structural analyses identify unusual quaternary structural rearrangements in the channel assembly which likely represent intermediate gating states. Moreover, a comparison with the published TRPM2_closed_ and TRPM2_open_ structures enabled us to speculate on the conformational pathway involving reduced symmetric rearrangements between the closed and open states of TRPM2.

## Results

### Structure determination and overall architecture of TRPM2_DR_

TRPM2_DR_ shares approximately 50% sequence identity with human TRPM2 (Supplementary Fig. 1), and when overexpressed in human embryonic kidney 293T (HEK293T) cells, wild-type TRPM2_DR_ channels exhibit large currents in response to direct application of ADPR and Ca^2+^ to the cytosolic side of inside-out patches (Supplementary Fig. 2). Notably, in the presence of saturating [Ca^2+^] (125 μM), TRPM2_DR_ shows an ADPR sensitivity similar to that of human TRPM2 ^23^.

For structural characterization, TRPM2_DR_ was purified in detergent in the presence of Ca^2+^ (TRPM2_DR_Ca2+_) or subsequently reconstituted into amphipol in the presence of both Ca^2+^ and ADPR (TRPM2_DR_ADPR/Ca2+_) (see methods). The TRPM2_DR_Ca2+_ and TRPM2_DR_ADPR/Ca2+_ structures were determined by cryo-EM to an overall resolution of ~3.8 Å and ~4.2 Å, respectively (Fig. 1 and Supplementary Figs. 3 and 4), enabling model building of ~85% of the TRPM2_DR_ polypeptide (see methods, Supplementary Fig. 5 and Table 1). In our study, we observed density corresponding to two unique structural features of the TRPM2 channel: a second coiled-coil and the NUDT9H domain at the C-terminus of the channel (Fig. 1). However, since these regions were the least well-defined in the EM maps, we facilitated model building and analysis by docking the homology model of the crystal structure of human ADP-ribose pyrophosphatase NUDT9 (PDB 1Q33) ^19^ into the NUDT9H density (see methods).

**Fig. 1:**
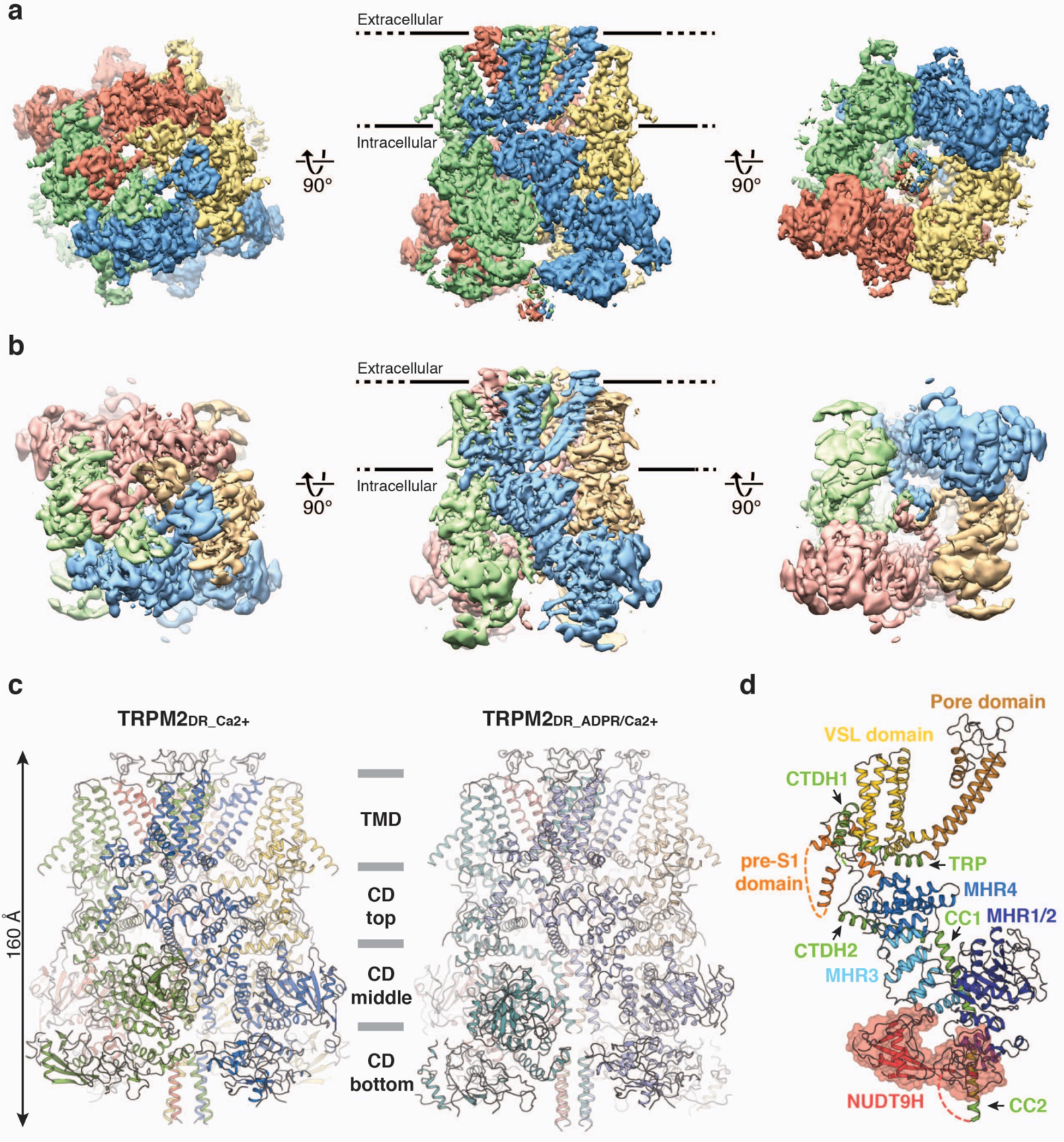
Overall architecture of TRPM2_DR_ structures. **a**,**b**, Cryo-EM reconstructions of TRPM2_DR_Ca2+_ (**a**) and TRPM2_DR_ADPR/Ca2+_ (**b**) structures viewed from the extracellular side (left), within the membrane bilayer (center), and from the cytosolic side (right). For TRPM2_DR_ADPR/Ca2+_, the NUDT9H domain is shown at 0.018 thresholding to emphasize conformational distinctions, while the rest of the molecule is shown at 0.033 thresholding. **c**, Cartoon representations of the TRPM2_DR_Ca2+_ (left) and TRPM2_DR_ADPR/Ca2+_ (right) structures with the same orientation as in the central panels of **a**and **b**. **d**, Detailed view of the green-colored protomer of TRPM2_DR_Ca2+_ structure in **c**. The NUDT9H domain is highlighted with a red surface representation. Dashed lines indicate loops and helices that were not modeled in the structure.

**Table 1.**
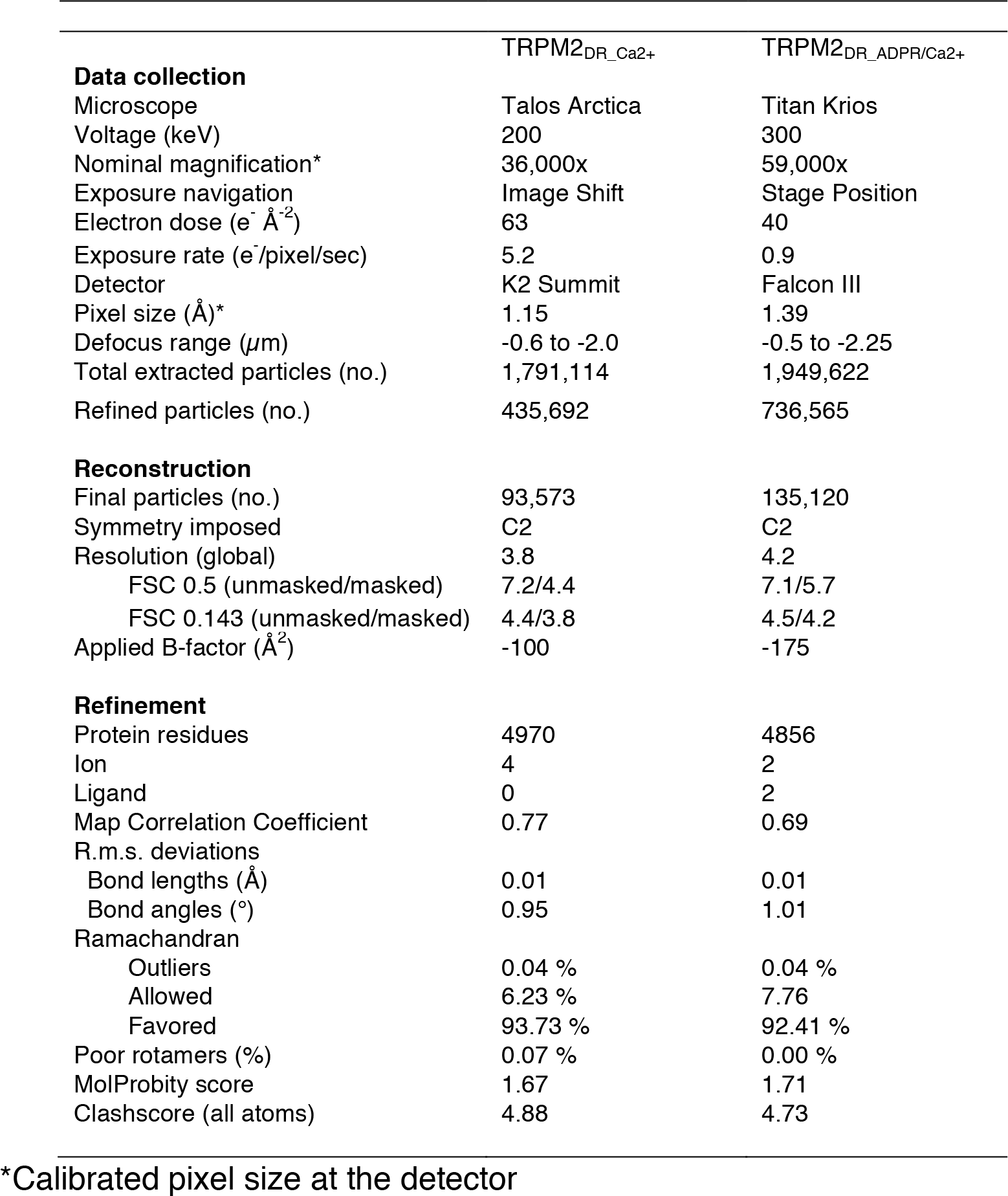
Data collection and refinement statistics.

Our TRPM2_DR_ channel forms a homo-tetramer (Fig. 1a,b). Viewed orthogonally to the membrane plane, the channel can be divided into four layers. Like other published TRP channel structures, the TMD layer adopts a domain-swapped configuration between the VSLD and the pore ^24,25^. Consistent with the previously reported TRPM2 structures ^8,26,27^, we also identified a Ca^2+^ binding site located in the cavity formed by the VSLD in the TRPM2_DR_Ca2+_ and TRPM2_DR_ADPR/Ca2+_ structures (Supplementary Fig. 6a-c). While the TMD and two membrane-proximal CD layers resemble the architecture observed in the TRPM8 and TRPM4 structures ^28–31^, the additional second coiled coil (CC2) and the NUDT9H domain together comprise a unique bottom CD layer in the TRPM2 structures (Fig. 1c). Similar to the previously reported TRPM2, TRPM4, and TRPM8 structures ^8,26–31^, each TRPM2_DR_ protomer contains an N-terminal region composed of MHR1 to MHR4, a transmembrane channel region, and a C-terminal region. The CC2 serves as a link between the C-terminal NUDT9H domain and the rest of the channel (Fig. 1d).

### TRPM2_DR_ structures in the intermediate states adopt two-fold symmetry

In contrast to the canonical four-fold symmetry reported in the TRPM2_closed_ and TRPM2_open_ structures, our TRPM2_DR_Ca2+_ and TRPM2_DR_ADPR/Ca2+_ structures adopt a two-fold symmetric arrangement when viewed along the central axis of the channel. This C2 symmetry was readily apparent upon reference-free 2D classification of the particle images (Supplementary Figs. 3e and 4e). To illustrate the key features associated with the observed two-fold symmetry, we focus first on the TRPM2_DR_Ca2+_ structure, which exhibits different magnitudes of two-fold symmetry across the layers of the channel. This reduced symmetry is most pronounced in the middle layer of the CD, comprising the MHR1/2 and MHR3 domains (Fig. 2a and Supplementary Fig. 7). Notably, comparisons with the published TRPM2_closed_ and TRPM2_open_ structures show that protomer A of our TRPM2_DR_Ca2+_ structure resembles the closed conformation while protomer B approximates the open conformation (Fig. 2b,c). This observation suggests that our TRPM2_DR_Ca2+_ structure may represent an intermediate state in which neighboring subunits adopt closed and the open conformations in an alternating manner, resulting in the two-fold symmetry between diagonally opposing subunits.

**Fig. 2:**
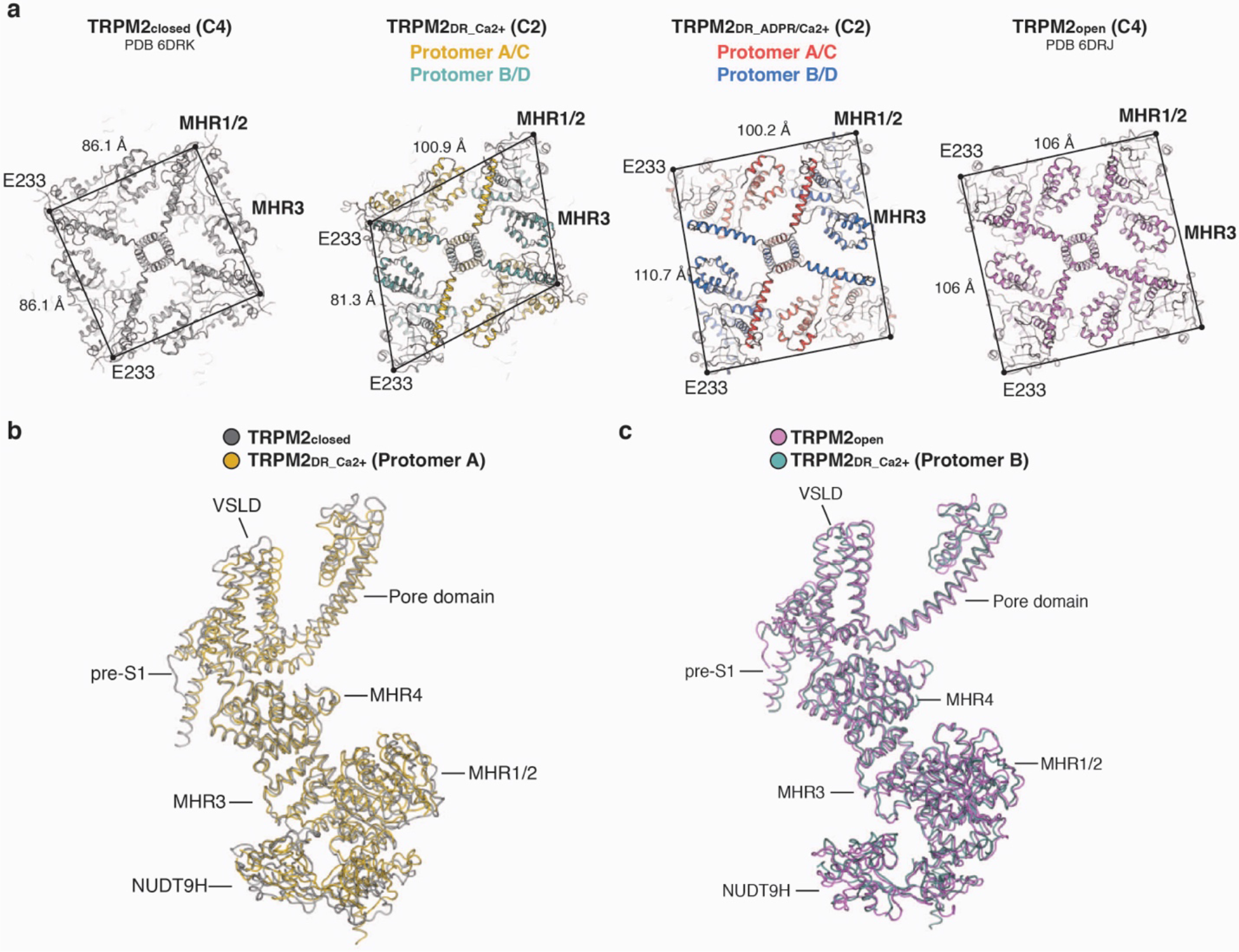
TRPM2_DR_ structures in the intermediate states. **a**, Extracellular views sliced through the middle layer of the CDs comprised of the MHR1/2 and MHR3 domains in the TRPM2_closed_, TRPM2_Ca2+_, TRPM2_ADPR/Ca2+_, and TRPM2_open_ structures. **b**,**c**, Structural alignment (cartoon representation) shows that protomer A in the TRPM2_DR_Ca2+_ structure (yellow) resembles the closed conformation in the TRPM2_closed_ structure (silver, PDB 6DRK) (**b**), while the protomer B (teal) resembles the open conformation in the TRPM2_open_ structure (violet, PDB 6DRJ) (**c**).

### Domain rearrangement at flexible junctions

To identify the origin of the two-fold symmetry, we performed global and local alignments of protomers A and B in the TRPM2_DR_Ca2+_ structure. The MHR3, MHR4, and VSLD are conformationally similar between protomers, whereas significant differences in domain arrangement are observed at three junctions: NUDT9H-MHR1/2, MHR1/2-MHR3, and VSLD-pore (Fig. 3a,b). When the MHR3 and MHR4 domains of protomers A and B are aligned, the MHR1/2 and NUDT9H domains exhibit a drastic rotation along individual axes (Fig. 3c). Moreover, this movement within the CD is propagated to the TM helices, resulting in two-fold symmetry within the TMD (Fig. 3d). The TMDs of the protomers A and B diverge at the S4b and the S4-S5 linker regions. In protomer A, the S4b adopts a 3_10_-helical structure, while the equivalent region in protomer B contains an α-helical structure and an unstructured loop. In addition, the S4-S5 linker in protomer A contains a π-helix, while the corresponding region in protomer B is α-helical and forms a continuous straight helix with S5. Due to the absence of a bend-inducing π-helix at the junction between the S4-S5 linker and S5, the entire pore domain of protomer B is positioned differently from that of protomer A with respect to the central axis of the channel. Therefore, the flexible elements in the S4b and the S4-S5 linker lead to a two-fold symmetric configuration of the TMD. Notably, a similar two-fold symmetric TMD arrangement induced by a conformational change in the S4-S5 linker was recently observed in the TRPV2 channel ^21^. The substantial conformational change at the S4b and S4-S5 linker between adjacent protomers leads to unexpectedly distinct configurations of the S6 gate. Although the distance between diagonally opposed gate residues in TRPM2_DR_Ca2+_ are larger than those observed in the TRPM2_closed_ structure, the S6 gate in TRPM2_DR_Ca2+_ is not as wide as that observed in the TRPM2_open_ structure (Supplementary Fig. 8a). Taken together, the structural differences between protomers A and B in TRPM2_DR_Ca2+_ originate at the flexible junctions located between NUDT9H-MHR1/2, MHR1/2-MHR3, and VSLD-pore. This flexibility allows the channel to assume a two-fold symmetric intermediate state, where two subunits approximate the closed conformation and the other two adopt an arrangement that is similar to the open state.

**Fig. 3:**
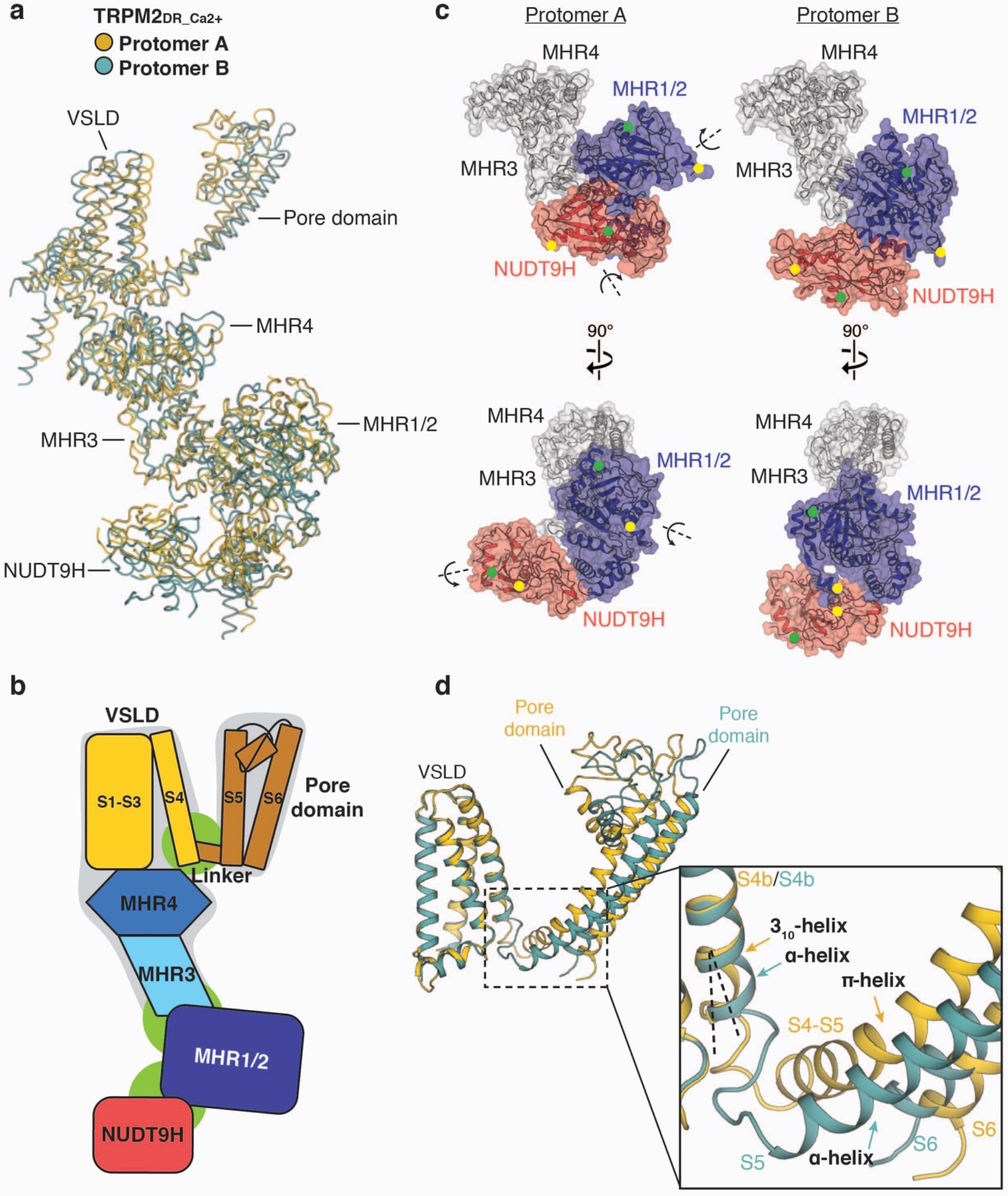
Flexible junctions enable conformational heterogeneity within the channel. **a**, Cartoon representation showing alignment between protomer A (yellow) and protomer B (teal) from the TRPM2_DR_Ca2+_ structure. **b**, Schematic diagram showing the three flexible junctions: NUDT9H-MHR1/2, MHR1/2-MHR3, and VSLD-pore (in green), and the static region: MHR3-MHR4-VSLD and the pore domain (in grey). The pre-S1 domain, TRP domain, and C-terminal helices are omitted for simplicity. **c**, Protomers A (left) and B (right) of the TRPM2_DR_Ca2+_ structure aligned at MHR3 and MHR4 (grey). Side-by-side surface representations indicate that the rotations of MHR1/2 and NUDT9H in protomer A around individual axes (left) lead to the orientations observed in protomer B (right). D90 (yellow) and Q137 (green) from MHR1/2 and V1372 (green) and H1468 (yellow) from NUDT9H are denoted as dots in both protomers. **d**, Protomers A (yellow) and B (teal) of the TRPM2_DR_Ca2+_ structure aligned at VSLD. Close-up view shows the structural divergence occurring at the S4b and the S4-S5 linker regions, giving rise to different configurations of the pore domain.

Importantly, the conformational differences between the closed and open states of TRPM2_DR_ are similar to those between two neighboring protomers in the TRPM2_DR_Ca2+_ structure (Supplementary Fig. 9). Given that protomers A and B in the TRPM2_DR_Ca2+_ structure resemble the closed and open conformation, respectively (Fig. 2b,c), we hypothesized that the addition of ADPR would induce further domain rearrangements at the junctions and enable transition of protomers A/C to the B/D configuration, converging the channel into a four-fold symmetric open conformation. To test this hypothesis, we prepared TRPM2_DR_ADPR/Ca2+_ by addition of ADPR to TRPM2_DR_Ca2+_ and determined its structure (Fig. 4). While protomer B in both structures retains the overall conformation similar to the open state (Fig. 4a), protomer A in TRPM2_DR_ADPR/Ca2+_ undergoes significant domain rearrangements in the CD at the interfaces between NUDT9H-MHR1/2 and MHR1/2-MHR3 (Fig. 4b). Notably, following the addition of ADPR, the CDs of protomers A and B in the TRPM2_DR_ADPR/Ca2+_ structure align well, which indicates that ligand binding changes the conformation of the CD in protomer A from the closed to the open state (Fig. 4c). Consequently, the CDs of the TRPM2_DR_ADPR/Ca2+_ structure converge to a nearly four-fold symmetric arrangement (Figs. 2a and 5).

**Fig. 4:**
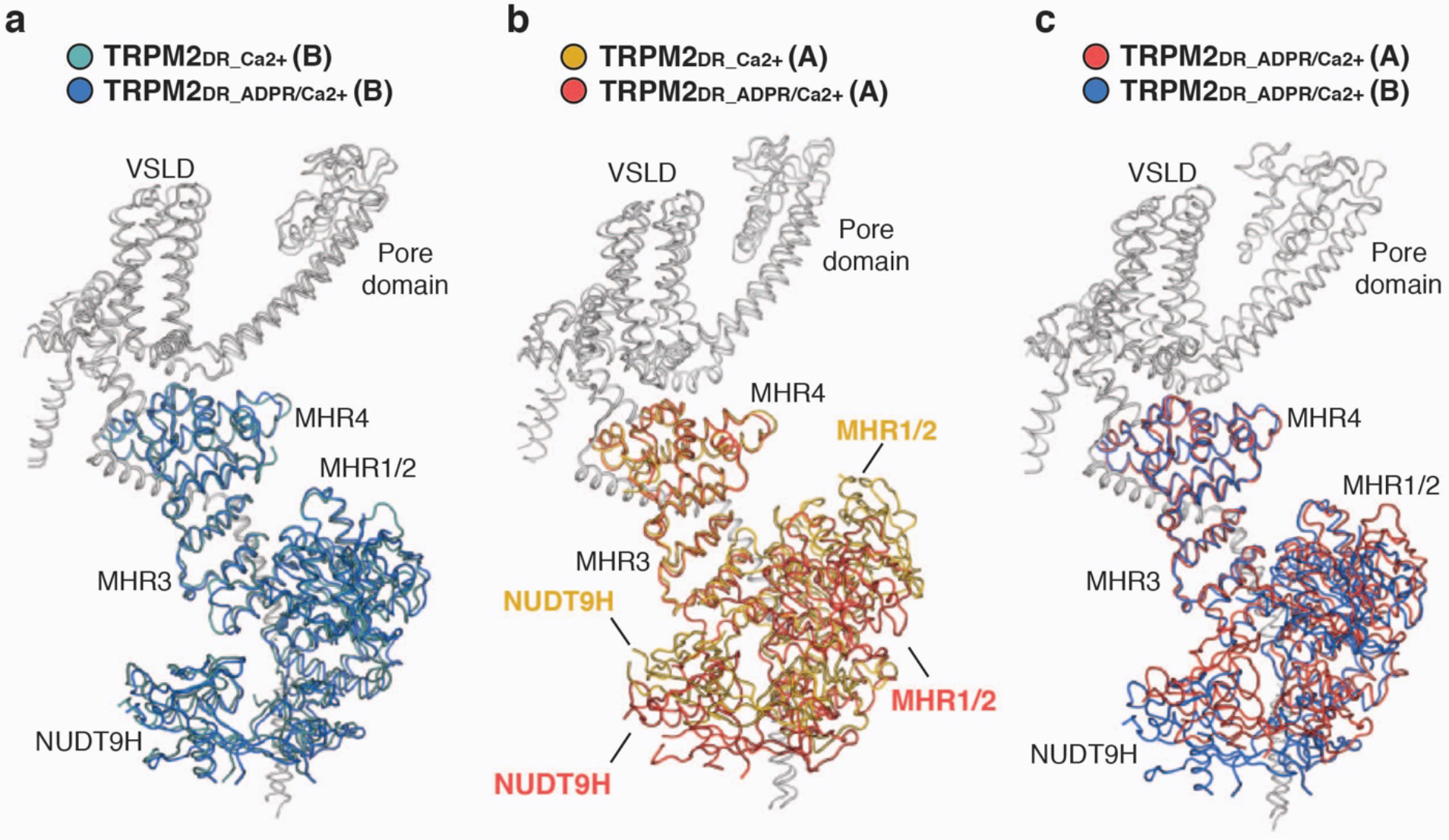
Addition of ADPR converts protomer configurations at the CD. **a-c**, Cartoon representations showing structural alignment of protomer B in the TRPM2_DR_Ca2+_ structure (teal) and protomer B in the TRPM2_DR_ADPR/Ca2+_ structure (blue) (**a**), protomer A in the TRPM2_DR_Ca2+_ structure (yellow) and protomer A in the TRPM2_DR_ADPR/Ca2+_ structure (red) (**b**), and protomer A (red) and protomer B (blue) in the TRPM2_DR_ADPR/Ca2+_ structure (**c**). Protomers are aligned at MHR4 domains. TMDs and C-terminal domains are colored in gray.

### Alternating quaternary structure rearrangement in the bottom CD layer

Based on these structural comparisons, we suggest that the four subunits of TRPM2 channel adopt a two-fold symmetric intermediate quaternary structure assembly before ultimately assuming the canonical four-fold symmetric arrangement in the open conformation. To visualize the conformational changes associated with reduced symmetry, we compared our structures with the published TRPM2_closed_ and TRPM2_open_ structures at the bottom layer of the CD, which is comprised of the mobile MHR1/2 and NUDT9H domains, and identified substantial rearrangements mediated by critical interfacial interactions (Fig. 5).

**Fig 5:**
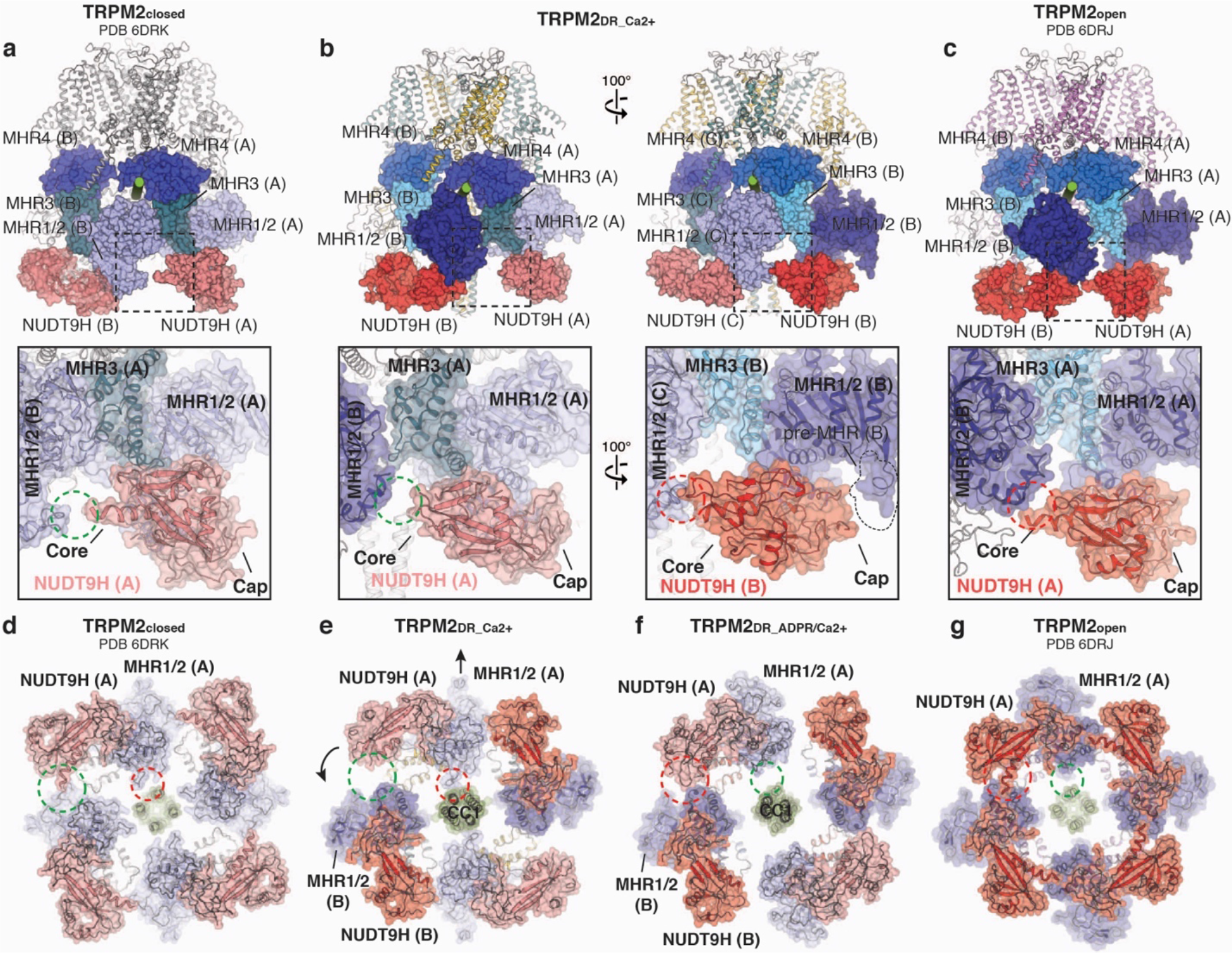
Alternating quaternary structure rearrangements in the cytoplasmic domain (CD) mediated by differential intra- and inter-subunit interactions. **a**–**c**, Comparison of the interaction networks between the NUDT9H domain and the neighboring MHR1/2 domain in the TRPM2_closed_ structure (**a**, PDB 6DRK), in the TRPM2_DR_Ca2+_ structure (**b**), and in the TRPM2_open_ structure (**c**, PDB 6DRJ). Close-up views show that the NUDT9H domain in the TRPM2_closed_ structure makes contact solely with MHR1/2 from the same subunit (a), while the Core subdomain of the NUDT9H in the TRPM2_open_ structure (**c**) makes additional interactions with the MHR1/2 from the adjacent protomer. More importantly, the TRPM2_DR_Ca2+_ structure adopts both interactions mediated by the NUDT9H and MHR1/2 domains (**b**). **d**–**g**, Viewed from the intracellular side, cartoon and transparent surface representations compare the conformational changes at the bottom layer of the CD in the TRPM2_closed_ structure (**d**), in TRPM2_DR_Ca2+_ (**e**) and TRPM2_DR_ADPR/Ca2+_ (**f**) structures from the current study, and in the TRPM2_open_ structure (**g**). Arrows indicate the domain movements observed in the TRPM2_DR_Ca2+_ (**e**) and TRPM2_DR_ADPR/Ca2+_ (**f**) structures relative to the channel in the closed conformation (**d**) and en route to the open conformation (**g**). Dashed circles highlight the detachment (green) and association (red) between the different domains as a result of structural rearrangements.

It was shown that in the apo conformation, the NUDT9H domain in the TRPM2_closed_ structure merely makes primary intra-subunit interactions with the MHR1/2 domain (Fig. 5a,d), while the binding of ADPR in the MHR1/2 domain relocates the NUDT9H domain to generate a secondary interface with the MHR1/2 domain from the neighboring protomer (Fig. 5c,g). Functional studies have shown that the interaction between the core subdomain of NUDT9H and the rest of the channel is important for stabilizing an open state of the TRPM2 channel, underscoring the importance of this secondary interface in channel opening ^20^. Strikingly, our TRPM2_DR_Ca2+_ structure adopts a combination of the two distinct interactions associated with the NUDT9H domain and the neighboring MHRs. Whereas the NUDT9H and MHR1/2 domains in protomer A exhibit only intra-subunit interactions, the core subdomain of the NUDT9H in protomer B makes additional contacts with the MHR1/2 domain in the neighboring protomer (Fig. 5b,e), resembling the interaction networks depicted in the TRPM2_closed_ and TRPM2_open_ structures, respectively. Furthermore, we observed changes in the interfacial interactions between the MHR1/2 domains and CC1. In the apo and closed state, the MHR1/2 domain is in loose contact with CC1 (Fig. 5d and Supplementary Fig. 10a). By contrast, in the open state, the binding of ADPR induces a displacement of the MHR1/2 domain, causing it to become disengaged from the CC1 helical bundle and positioned away from the central axis (Fig. 5g and Supplementary Fig. 10d). Our TRPM2_DR_Ca2+_ structure adopts both types of interactions between the MHR1/2 domain and CC1, further indicating a hybrid of the closed and the open conformations in the CD (Fig. 5e).

Therefore, based on the comparison with the TRPM2_closed_ and TRPM2_open_ structures, the alternating quaternary organization of the CDs in the TRPM2_DR_Ca2+_ structure suggests that ADPR-binding might induce further rearrangements that ultimately converge the reduced symmetry to adopt the four-fold symmetry apparent in the TRPM2_open_ structure. In our TRPM2_DR_ADPR/Ca2+_ structure, we observe density for ADPR in the cleft of the MHR1/2 domain in protomers B and D, the shape and location of which resemble those depicted in the TRPM2_open_ structure in complex with Ca^2+^ and ADPR (Supplementary Fig. 6e). Although it is difficult to identify corresponding ADPR density in protomers A and C due to the low resolution of the NUDT9H and MHR1/2 domains, we observed that the addition of ADPR converts protomers A/C from the closed to the open conformation, wherein the MHR1/2 domains detach from CC1 and the NUDT9H domains form interactions with neighboring MHR1/2 domains in all four protomers (Fig. 5f). As a result, the CDs in the TRPM2_DR_ADPR/Ca2+_ structure have been converted to a nearly four-fold symmetric arrangement that is similar to the CD conformation shown in the TRPM2_open_ structure.

While its CDs adopt a nearly four-fold symmetric conformation, the TMDs of our TRPM2_DR_ADPR/Ca2+_ structure still adopt a hybrid of the closed and the open conformations (Supplementary Fig. 8b-e), indicating that the TMDs have not converged towards the open conformation. This suggests that the captured TRPM2_DR_ADPR/Ca2+_ structure may represent an intermediate state in which the structural rearrangements occurring at the CD have not been fully propagated to the TMD.

### Subunit-subunit interfaces in the middle CD layer

The interfaces between the MHR1/2 and MHR3 domains in adjacent subunits comprise the middle layer of the CD, which undergoes the largest conformational changes between the closed and the open state upon ADPR binding. Notably, the reduced symmetry is most pronounced in the middle layer of the CD in the TRPM2_DR_Ca2+_ structure (Fig. 2a). Comparison between the published TRPM2_DR_closed_ and TRPM2_DR_open_ structures showed that ADPR-induced conformational changes between MHR1/2 and MHR3 in each protomer leads to drastic changes in the subunit-subunit interface in the middle CD layer (Fig. 6a,c). While the closed state of TRPM2_DR_ exhibits large interfacial interactions between neighboring MHR1/2 and MHR3 domains, the corresponding areas are significantly smaller in the open state (Fig. 6d,f). Notably, in the TRPM2_DR_Ca2+_ structure, the middle CD layer accommodates two distinct subunit-subunit interfaces (Fig. 6b,e), consistent with the most pronounced two-fold symmetry observed in this region (Fig. 2a). While the interface between MHR1/2 in protomer A and the neighboring MHR3 is similar to the interfacial network in the closed state, the contact mediated by MHR1/2 in protomer B and the adjacent MHR3 domain resembles that of the open state. Our observation of these alternating interfaces in the TRPM2_DR_Ca2+_ structure prompted us to question why such a two-fold symmetric intermediate state might exist. We posit that concerted quaternary structural changes from the closed to the open state would invoke substantial rearrangements at the subunit-subunit interfaces, which could be energetically costly. However, by adopting an intermediate state with alternating subunit-subunit interfaces, the transition from the closed to the open state through a two-fold symmetric intermediate could reduce the energetic barrier for each structural rearrangement, thereby facilitating the channel gating (Fig. 6).

**Fig. 6:**
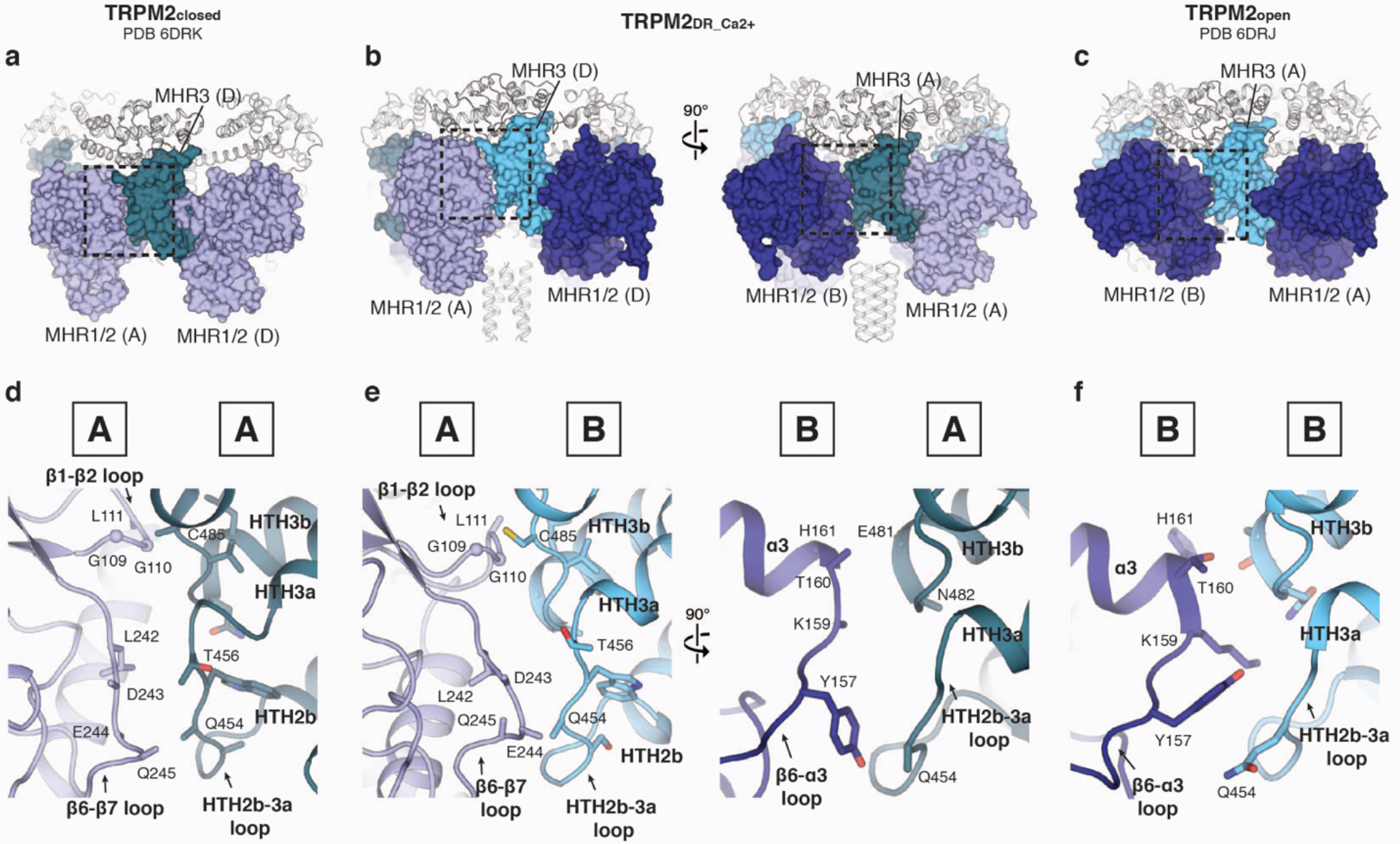
Subunit-subunit interfaces in the middle layer of CD. **a-c**, Comparison of the interaction networks between the neighboring subunits mediated by MHR1/2 and MHR3 domains in the TRPM2_closed_ structure (**a**, PDB 6DRK), in the TRPM2_DR_Ca2+_ structure (**b**), and in the TRPM2_open_ structure (**c**, PDB 6DRJ). **d**–**f**, Close-up views of regions highlighted by dashed squares in **a**–**c** correspondingly. In the TRPM2_DR_Ca2+_ structure, the interface between MHR1/2 in protomer A and MHR3 in protomer D (equivalent to B) (**e**, left panel) resembles the interfacial network in the TRPM2_closed_ structure (**d**); the interface between MHR1/2 in protomer B and MHR3 in protomer A (**e**, right panel) is similar to that in the TRPM2_open_ structure (**f**).

## Discussion

In this study, we observe a two-fold symmetric arrangement in the homo-tetrameric channel TRPM2 in complex with only Ca^2+^ (TRPM2_DR_Ca2+_), and with both ADPR and Ca^2+^ (TRPM2_DR_ADPR/Ca2+_), a feature previously observed in the TRPV2 and TRPV3 channels ^21,22^. Interestingly, comparing our structures with the recently published four-fold symmetric TRPM2_closed_ and TRPM2_open_ structures reveals that protomers A and C of TRPM2_DR_Ca2+_ resemble the closed conformation, while protomers B and D approximate the open conformation, indicating that the homo-tetramer can accommodate a hybrid of structural arrangements that are representative of the closed and the open states.

We consider the possible reason for the unique two-fold symmetric arrangement observed in our TRPM2_DR_ structures may lie in the biochemical preparation of our samples. While the construct (full-length wild type TRPM2_DR_), expression system (HEK293 cells), and detergent (digitonin) we used were very similar to those of the previously published structures ^8^, our biochemical preparation also included Ca^2+^. Notably, our TRPM2_DR_Ca2+_ structure is the only structure of TRPM2 to date that was determined in the presence of Ca^2+^ only, thus enabling us to dissect the role of Ca^2+^ in TRPM2 gating. It is possible that Ca^2+^ binding in the VSLD confers flexibility to the junction at the S4-S5 linker, thus priming the channel for opening. Consistent with this idea, the density for Ca^2+^ in the TRPM2_DR_Ca2+_ reconstruction is stronger in protomer B compared to that of protomer A (10.8 σ versus 9.5 σ, respectively) (Supplementary Fig. 6a,b). In addition, we cannot exclude another possibility that endogenous ADPR was captured in the Ca^2+^-bound TRPM2 during sample preparation, as we also observe weak densities in the cleft of the MHR1/2 domain in protomers B and D of the TRPM2_DR_Ca2+_ reconstruction, which may correspond to ADPR (Supplementary Fig. 6d). However, the quality of the density in these regions was insufficient to unambiguously assign these detailed structural components.

Taken together, the alternating closed and open conformations of neighboring protomers observed in both our TRPM2 structures, as well as the conversion of the CD to a four-fold symmetric open conformation in the presence of additional ADPR, suggests that the TRPM2 channel adopts two-fold symmetric intermediate states en route to the open state (Fig. 7c). Reduced symmetry transitions could potentially be a stepwise mechanism used to accommodate the substantial conformational changes that occur during channel activation. Consistent with this idea, our interface analysis suggests that the two-fold symmetric intermediates confer an advantage by reducing the energetic barrier to the concerted quaternary structural changes going from the closed to the open state (Figure 6). A similar departure from the canonical four-fold symmetry associated with TRP channel gating was recently observed during ligand-induced gating in the TRPV2 and TRPV3 channels ^21,22^, suggesting that adoption of a C2-symmetric conformation may be a more widely used mechanism in the TRP channel superfamily.

**Fig. 7:**
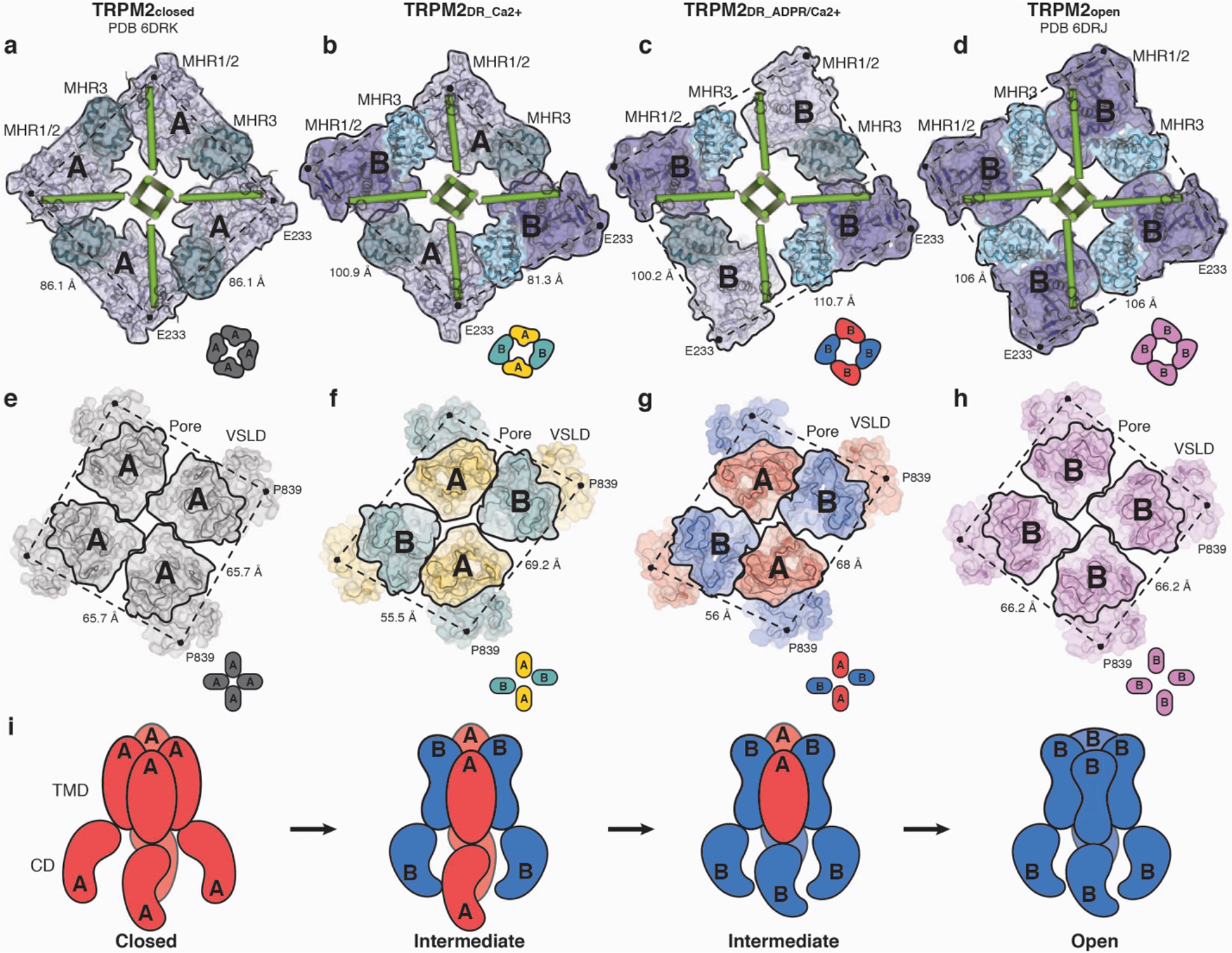
A trajectory of conformational changes during in the TRPM2_DR_ channel gating. **a**–**d**, Viewed from the extracellular side, surface and cartoon representations of the middle layer of CD sliced through TRPM2_closed_ (**a**), TRPM2_DR_Ca2+_ (**b**), TRPM2_DR_ADPR/Ca2+_ (**c**), and TRPM2_open_ (**d**) structures. Individual protomers are highlighted in black frames. The protomer configurations are indicated by type “A” or “B” and also depicted as cartoon diagrams in insets. Cα atoms of the E233 residues are shown as black dots, and distances between Cα atoms are indicated (Å). **e**–**h**, Viewed from the extracellular side, surface and cartoon representations of the TMD of TRPM2_closed_ (**e**), TRPM2_DR_Ca2+_ (**f**), TRPM2_DR_ADPR/Ca2+_ (**g**), and TRPM2_open_ (**h**) structures. Individual pore domains are highlighted in black frames. The protomer configurations are indicated by type “A” or “B” and also depicted as cartoon diagrams in insets. Cα atoms of the P839 residues are shown as black dots, and distances between Cα atoms are indicated (Å). **i,**Cartoon diagram of alternating quaternary structural rearrangements in TRPM2 channel gating.

## Acknowledgments

Cryo-EM data for TRPM2_DR_Ca2+_ and TRPM2_DR_ADPR/Ca2+_ were collected at The Scripps Research Institute (TSRI) electron microscopy facility, and the Duke Shared Materials Instrumentation Facility (SMIF), respectively. Preliminary cryo-EM work, including sample screening, was performed at the cryo-EM facility at NIEHS as part of the Molecular Microscopy Consortium (MMC) in the Research Triangle, North Carolina. We thank J.C. Ducom at the TSRI High Performance Computing facility for computational support and B. Anderson for microscope support. We thank L. Zubcevic, A. Bartesaghi, L. Csanady for providing critical manuscript reading, a preprocessing interface, and preliminary functional studies, respectively. This work was supported by the National Institutes of Health (R35NS097241 to S.-Y.L., DP2EB020402 and R21AR072910 to G.C.L.) and by the National Institute of Health Intramural Research Program; US National Institutes of Environmental Health Science (NIEHS) (ZIC ES103326 to M.J.B). G.C.L is supported as a Searle Scholar, a Pew Scholar in the Biomedical Sciences, supported by the Pew Charitable Trusts, and by an Amgen Young Investigator award. M.W is supported by a National Science Foundation Graduate Student Research Fellowship. Computational analyses of EM data were performed using shared instrumentation funded by NIH S10OD021634.

## Author Contributions

Y.Y. conducted all biochemical preparation, cryo-EM experiments and single-particle 3D reconstruction of TRPM2_DR_ADPR/Ca2+_, and model building and refinement under the guidance of S.-Y.L. M.W. conducted cryo-EM experiments and single-particle 3D reconstruction of TRPM2_DR_Ca2+_ under the guidance of G.C.L. A.H. collected cryo-EM data of TRPM2_DR_ADPR/Ca2+_, and helped with cryo-EM sample screening under the guidance of M. Borgnia. W.F. carried out electrophysiological recordings under the guidance of S.-Y.L. Y.Y. S.-Y.L., G.C.L, and M.W. wrote the paper.

## Declaration of Interests

The authors declare no competing interests.

## Methods

### Protein expression and purification

Zebrafish TRPM2 showed optimal biochemical stability based on a screen of eight orthologues including human and rat TRPM2. A codon-optimized and full-length gene for zebrafish (*Danio rerio*, XP_009303266.1) TRPM2 (TRPM2_DR_) was synthesized and cloned into a modified pEG BacMam vector ^32^ in frame with a C-terminal FLAG affinity tag (Bio Basic Inc.). The full length wild-type protein was expressed by baculovirus-mediated transduction of HEK293S GnTI^−^ (*N*-acetylglucosaminyltransferase I-negative) suspension cells. In brief, the wild-type construct was transformed into DH10Bac *Escherichia coli* cells and the recombinant bacmid was isolated and transfected into *Spodoptera frugiperda* (Sf9) cells. The BacMam virus was generated, amplified and added to HEK293S GnTI^−^ cells (ATCC) at a cell density of 3-3.5 M ml^−1^. Cells were grown in Freestyle 293 media (Life Technologies) supplemented with 2% (v/v) FBS(Gibco) at 37°C in the presence of 8% CO_2_. After 18 h, 10 mM sodium butyrate was added to the cell culture and the temperature was reduced to 30 °C to boost protein expression. 43-48 h after induction, cells were harvested and resuspended in buffer A (50 mM Tris pH 8, 150 mM NaCl, 1% digitonin (Sigma-Aldrich), 6.4 mM β-mercaptoethanol, 12 μg mL^−1^ leupeptin, 12 μg mL^−1^ pepstatin, 12 μg mL^−1^ aprotinin, 1.2 mM phenylmethylsulfonyl fluoride and DNase I). Phytic acid (InsP_6_, Sigma-Aldrich) and CaCl_2_ were added to a final concentration of 1 mM and 2 mM respectively. The protein was solubilized at 4 °C for 1 h. The insoluble material was removed by centrifugation at 8,000*g* for 30 min. The supernatant was incubated with anti-FLAG M2 resin (Sigma-Aldrich) for 30-40 min at 4 °C with gentle agitation. The resin was washed with ten column volumes of filtered buffer B (20 mM Tris pH 8, 150 mM NaCl, 0.06-0.1 % digitonin, 1 mM InsP_6_, 2 mM CaCl_2_ and 2 mM DTT) and eluted with five column volumes of the same buffer supplemented with 0.1 mg mL^−1^ FLAG peptide. The elution was concentrated and further purified on a Superose 6 10/300 GL column (GE Healthcare) equilibrated with filtered buffer B. The peak fractions were concentrated to 0.5 mg mL^−1^ for cryo-EM analysis of the TRPM2_DR_Ca2+_ structure.

To solve the TRPM2_DR_ADPR/Ca2+_ structure in complex with both ADPR and Ca^2+^, after purifying the protein by size-exclusion chromatography as described above, peak fractions were combined and mixed with Amphipol PMAL-C8 (Avanti polar lipid) at 1:10 (wt/wt) ratio and incubated overnight at 4°C with gentle agitation. To remove detergent, 25 mg ml^−1^ Bio-Beads SM-2 (Bio-Rad) was added and incubated at 4°C for 1 h. After removing the Bio-Beads, the reconstituted protein was mixed with 50 μM ADP-Ribose (Sigma-Aldrich) and further purified on a Superose 6 10/300 GL column (GE Healthcare) equilibrated with buffer C (20 mM Tris pH8, 150 mM NaCl, 0.5 mM CaCl_2_, 50 μM ADPR). The peak fractions were collected and concentrated to ~1.25 mg mL^−1^ and incubated with ADPR to a final concentration of ~1 mM for 30-45 min before sample freezing for cryo-EM study.

### Cryo-EM sample preparation

For cryo-EM study of the TRPM2_DR_Ca2+_ structure, a thin amorphous carbon film was floated onto UltrAuFoil^®^ R1.2/1.3 300-mesh grids (Quantifoil). 3 μL of TRPM2 (0.5 mg/ml) was applied to freshly plasma cleaned grids, and the grids were manually blotted ^33^ using a custom-built manual plunger in a cold room (≥95% relative humidity, 4 °C). Sample was blotted for ~4 s with Whatman No.1 filter paper immediately prior to plunge freezing in liquid ethane cooled by liquid nitrogen.

The cryo-EM grids for the analysis of the TRPM2_DR_ADPR/Ca2+_ structure were prepared by applying 3 μL PMAL-C8 reconstituted TRPM2 (~1.25 mg mL^−1^) in the presence of ADPR and Ca^2+^ to freshly glow-discharged UltrAuFoil^®^ R1.2/1.3 300-mesh grids (Quantifoil). The grids were blotted for 2 s at 25 °C under 95% humidity before being plunged into liquid ethane using a Leica EM GP2 (Leica Microsystems). The grids were stored in liquid nitrogen before data acquisition.

### Cryo-EM data acquisition and data processing

#### TRPM2_DR_Ca2+_ structure

The Cryo-EM data used for the TRPM2_DR_Ca2+_ structure were acquired using the Leginon automated data acquisition program ^34^. All image pre-processing (frame alignment, CTF estimation, particle picking) were performed in real-time using the Appion image processing pipeline ^35^ during data collection. Images of frozen hydrated TRPM2 were collected on a Talos Arctica (Thermo Fisher) TEM operating at 200 keV. Movies were collected using a K2 Summit direct electron detector (Gatan) in counting mode at a nominal magnification of 36,000x corresponding to a physical pixel size of 1.15 Å/pixel. A total of 3,039 movies (64 frames/movie) of TRPM2 were collected by navigating to the center of four holes and image shifting ~2 μm to each exposure target. Movies were collected using a 16 second exposure with an exposure rate of 5.2 e^−^/pixel/sec, resulting in a total exposure of ~63 e^−^/Å^2^ (1.17 e^−^/Å^2^/frame) and a nominal defocus range from −1.2 μm to −2 μm. The MotionCor2 frame alignment program ^36^ was used to perform motion correction and dose-weighting as part of the Appion pre-processing workflow. Frame alignment was performed on 5 × 5 tiled frames with a B-factor of 100 applied. Unweighted summed images were used for CTF determination using CTFFIND4 ^37^. Difference of Gaussians (DoG) picker ^38^ was used to automatically pick particles from the first 636 dose-weighted micrographs yielding a stack of 176,961 particles that were binned 4 × 4 (4.6 Å/pixel, 80 pixel box size) and subjected to reference-free 2D classification using RELION 2.1 ^**39**^. The best nine classes were then used for template-based automated particle picking against the whole dataset using RELION. A total of 1,791,114 particles were extracted from these micrographs and binned 4 × 4 (4.6 Å/pixel, 80 pixel box size). Reference-free 2D classification in RELION was then used to sort out non-particles and poor-quality picks in the data. A total of 435,692 particles corresponding to 2D class averages that displayed strong secondary-structural elements were input to 3D auto-refinement in RELION without symmetry imposed. EMD-7127 was low-pass filtered to 30 Å and used as an initial model. The refined particle coordinates were then used for re-centering and re-extraction of particles binned 2 × 2 (2.3 Å/pixel, 160 pixel box size). The resulting stack was subjected to 3D auto-refinement using the map obtained from the previous refinement as an initial model, and with a soft mask (5 pixel extension, 5 pixel soft cosine edge) generated from a volume contoured to display the full density. These particles were then subjected to 3D classification (k=6, tau fudge=12) without angular or translational searches using the same soft mask. Particles contributing to the classes that possessed the best resolved densities around the transmembrane domain of the channel were 3D auto-refined and then re-centered and re-extracted without binning (135,215 particles, 1.15 Å/pixel, 320 pixel box size). These particles were then 3D auto-refined and subjected to 3D classification (k=3, tau fudge=12) without angular or translational searches. Two classes, comprising 94,028 particles, displayed the best-resolved density around the transmembrane region and C2 symmetry. 3D auto-refinement of these particles with C2 symmetry imposed yielded a ~3.9 Å reconstruction as determined by gold-standard 0.143 Fourier shell correlation (FSC) ^40^, using phase-randomization to account for the convolution effects of a solvent mask on the FSC between the two independently refined half maps ^41^. To improve map quality, all particles collected at greater than 2 μm defocus were removed. 3D auto-refinement of the final particle stack (93,573 particles) with C2 symmetry imposed yielded a ~3.8 Å reconstruction as determined by gold-standard 0.143 FSC.

#### TRPM2_DR_ADPR/Ca2+_ structure

For the TRPM2_DR_ADPR/Ca2+_ structure, the cryo-EM data were acquired using the EPU automated data-acquisition program. Images were collected on a Titan Krios (FEI) operating at 300 keV equipped with a Falcon III direct electron detector operating in counting mode. 2496 movies were collected at a nominal magnification of 59,000x with a physical pixel size of 1.39 Å/pixel using a nominal defocus range of −0.5 to −2.25 μm. Each movie (45 frames) was acquired using a dose rate of ~0.91 e^−^/pixel/s and a total exposure of ~40 e^−^/Å^2^.

Motion correction and dose-weighting was performed using the MotionCor2 frame alignment program on 5 × 5 tiled frames with a B-factor of 150 applied ^36^. Gctf ^42^ was used for CTF estimation of unweighted summed images and 2426 good micrographs were selected. 1719 particles were manually picked and subject to reference-free 2D classification (k=10, tau=2) in RELION 3.0 ^43^. The best seven classes were used as templates for auto-picking of a total of 1,949,622 particles. The dataset was extracted unbinned and subjected to reference-free 2D classification, construction of ab-initio model, heterogeneous refinement, and homogeneous refinement in CryoSPARC ^44^, yielding a 3D reconstruction of ~4.3 Å resolution. In parallel, the 1,949,622 particles were extracted, Fourier binned 4 × 4 (5.56 Å/pixel, 64 pixel box size) and subjected to reference-free 2D classification. Good 2D classes showing secondary structural features were selected. After further removing micrographs with a figure of merit (FOM) below 0.1 and with astigmatism above 500 nm, a total of 736,565 particles from 2366 good micrographs were combined and input to 3D auto-refinement in RELION with C1 symmetry. The model of ~4.3 Å resolution generated by CryoSPARC was low-pass filtered to 30 Å and used as an initial model without a reference mask. The refined particles were re-centered, re-extracted, Fourier binned 2 × 2 (2.78 Å/pixel, 128 pixel box size), and subject to 3D auto-refinement with C1 symmetry, using the model from the previous auto-refinement as the reference mask and with a soft mask (5 pixel extension, 5 pixel soft cosine edge). The refined particles were input to 3D classification (k=4, tau=12) without alignment using the same mask. 135,120 particles comprising the best 3D class which shows the most well-defined cytoplasmic domain (CD) were re-centered, re-extracted, unbinned, and subject to 3D auto-refinement with C2 symmetry with a mask around the full density, yielding a final construction of ~4.2 Å resolution determined by gold-standard 0.143 Fourier shell correlation (FSC) using RELION 3.0 ^43^. Per-particle CTF refinement and Bayesian polishing ^45^ were attempted, but the resolution was not improved.

### Model building and refinement

An initial model of TRPM2 was generated by the RaptorX structure prediction server ^46^ with the sequence of zebrafish TRPM2 (XP_009303266.1). For the model building of the TRPM2_Ca2+_ structure, individual domains (NUDT9H, VSLD, pre-S1, pore, CC1, CC2, MHR1/2, MHR3, and MHR4) were manually docked into the electron density map and the subsequent model building was performed manually in Coot ^47^. A series of rigid-body fitting was performed for secondary structures within the structural domains, including the MHR domains, the pre-S1 domain, VSLD, the pore domain, and the NUDT9H domain. Residues with bulky side chains guided the correct register of helices and β-strands. Side chains were adjusted to optimal rotamer conformations and some loops that connect secondary structures were rebuilt to fit in the electron density. Unstructured loops and side chains whose electron density was not resolved in the map were deleted from the model. To facilitate the model building of the NUDT9H domain, the crystal structure of human ADP-ribose pyrophosphatase NUDT9 (PDB ID: 1Q33) was docked into the electron density. The coordinates of the TRPM2_DR_Ca2+_ structure was docked into cryo-EM map for the TRPM2_ADPR/Ca2+_ structure, and the subsequent model building was performed in a similar manner in Coot. Ideal geometry restraints were imposed on the secondary structure and the rotamer conformation as much as possible during the initial manual model building in Coot. In the TRPM2 _DR_Ca2+_ structure, a calcium ion was placed into the EM density at the putative binding site in each protomer; while in the TRPM2 _DR_ADPR/Ca2+_ structure, a calcium ion and an ADPR molecule were built into the putative EM density in protomers B and D. The ligand restraints were generated by eLBOW ^48^. The two structure models were subsequently real-space refined in the PHENIX graphical interface against the cryo-EM map along with ligand restraints, using global minimization and rigid body refinement with secondary structure restraints ^49^. Problematic regions in the real-space refined structure were identified using the Molprobity server (http://molprobity.biochem.duke.edu/) ^50^ and were manually fixed in Coot. FSCs of the half maps against the refined model agree with each other, indicating that the model is not over-refined (Supplementary Figs. 3 and 4). The final TRPM2_DR_ structures cover about ~85% of the entire sequence. The TRPM2 _DR_Ca2+_ structure model consists of amino acids 40-1467 for chains A and C and 40-1470 for chains B and D with a few connections missing (1-39, 53-88, 518-534, 562-588, 764-797, 1115-1119, 1185-1187, 1210-1245, 1291-1302, 1338-1340, 1350-1354, 1364-1368, 1384-1391, 1425-1426, 1436-1443, and 1468-1474 in chains A and C; 1-39, 53-87, 292-295, 518-525, 561-588, 766-796, 865-868, 1115-1119, 1185-1187, 1210-1246, 1293-1302, 1367-1368, 1382-1385, 1420-1423, and 1471-1474 in chains B and D). The TRPM2 _DR_ADPR/Ca2+_ structure model contains amino acids 98-1470 for chains A and C and 40-1470 for chains B and D with several regions missing (1-97, 292-295, 518-534, 559-590, 762-797, 880, 1113-1119, 1185-1187, 1210-1251, 1292-1302, 1367-1368, 1382-1385, 1420-1423, 1471-1474 for chains A and C, while 1-39, 52-94, 109-113, 290-296, 518-534, 560-589, 766-797, 865-868, 1112-1119, 1185-1187, 1210-1247, 1293-1302, 1367-1368, 1382-1385, 1420-1423, 1438-1441, 1471-1474 for chains B and D). Polyalanine models were assigned for the second coiled coil (1188-1209) in both structures.

### Electrophysiology

HEK293T cells (62312975 – ATCC) were grown in DMEM supplemented with 10% FBS (Gibco), 1% penicillin/streptomycin (Gibco) and were sustained in 5% CO_2_ atmosphere at 37ºC. Cells between passage 10 – 30 grown in 40-mm wells were transiently transfected at 30 – 50% confluency with plasmids encoding for TRPM2_DR_ and green fluorescent protein (GFP) using FuGene6 (Promega). Transfected cells were subcultured onto laminin (Sigma-Aldrich) coated 12-mm round glass coverslips (Fisher Scientific) 24 hours post transfected and used 12 – 24 hours after for electrophysiological measurements.

Voltage-clamp recordings were performed in the inside-out patch clamp configuration with glass electrodes pulled from borosilicate glass capillaries (Sutter Instruments) with a final resistance of 1.5 – 2.5 MΩ. The extracellular pipette solution contained (in mM) 140 NaCl, 5 KCl, 2 MgCl_2_, 10 HEPES, 5 EGTA, and adjusted to pH 7.4 (NaOH). Glass coverslips were placed in an open bath chamber (RC-26G, Warner Instruments) filled with the external pipette solution and after obtaining a GΩ seal, the patch was excised into the bath. An intracellular bath solution containing (in mM) 140 NaCl, 5 KCl, 2 MgCl_2_, 10 mM HEPES, 1 EGTA, 1.12 CaCl_2_ at pH 7.4 (NaOH), with estimated free [Ca^2+^] ≈125 μM calculated with MaxChelator software (http://maxchelator.stanford.edu) ^51^ with and without varying concentrations of ADPR (Sigma) (prepared daily from aqueous stock solutions (50 mM) stored at −80º C) was applied to the inside of the patch with a pressurized perfusion system.

Inside-out current responses were elicited with a continuous repeating voltage ramp protocol (holding potential 0 mV for 50-ms before and after a 400-ms voltage ramp from 0 to +50 mV). Current responses were low-pass filtered at 1 – 2 kHz (Axopatch 200B), digitally sampled at 5 – 10 kHz (Digidata 1440A), converted to digital files in Clampex10.7 (Molecular Devices) and stored on an external hard drive for offline analyses (Clampfit10.7, Molecular Devices; OriginPro 2016, OrginLab Corp.). The inward current at +50 mV (V_m_ = −50 mV) from each patch was used to calculate the mean current amplitude for each ADPR concentration.

### Data availability statement

The sequence of TRPM2_DR_ can be found in the National Center for Biotechnology Information under accession code XP_009303266.1. For the TRPM2_DR_Ca2+_ and TRPM2_DR_ADPR/Ca2+_ structures, the coordinates have been deposited in the Protein Data Bank with the PDB ID ### and ### and the cryo-EM density maps have been deposited in the Electron Microscopy Data Bank with the accession number EMD-### and EMD-###.

---

**Supplementary Figure 1.**
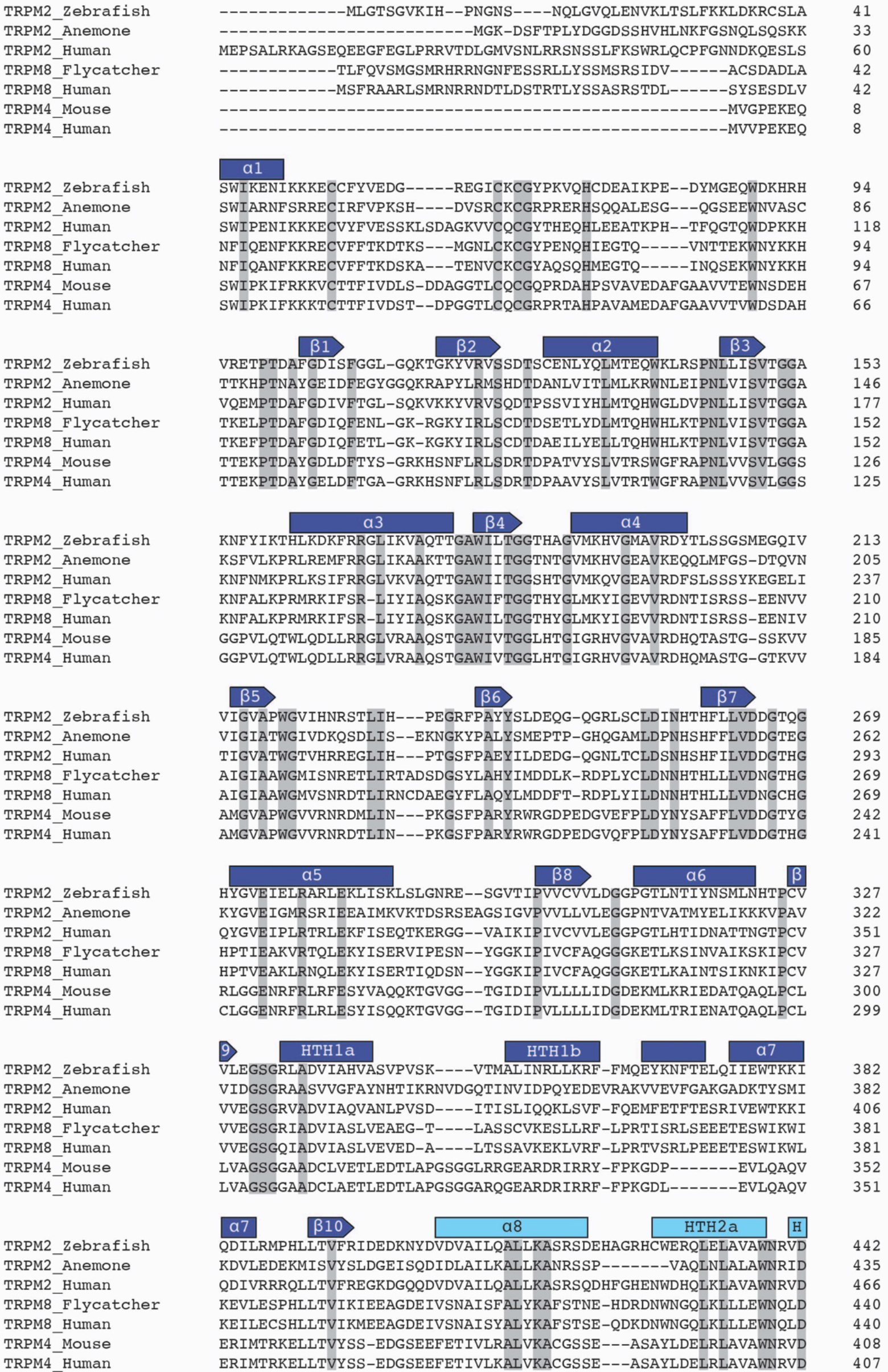
Sequence alignment of TRPM2 with representative TRPM8 and TRPM4 orthologues. Secondary structures in the TRPM2DR structure are indicated by rectangles (helices) and arrows (β-strands) and are colored as in Fig. 1d. Residues with absolute conservation are highlighted in grey. Sequences corresponding to the NUDT9H domain in zebrafish and human TRPM2 channels are indicated by red bars. Green line identifies the Nudix box.

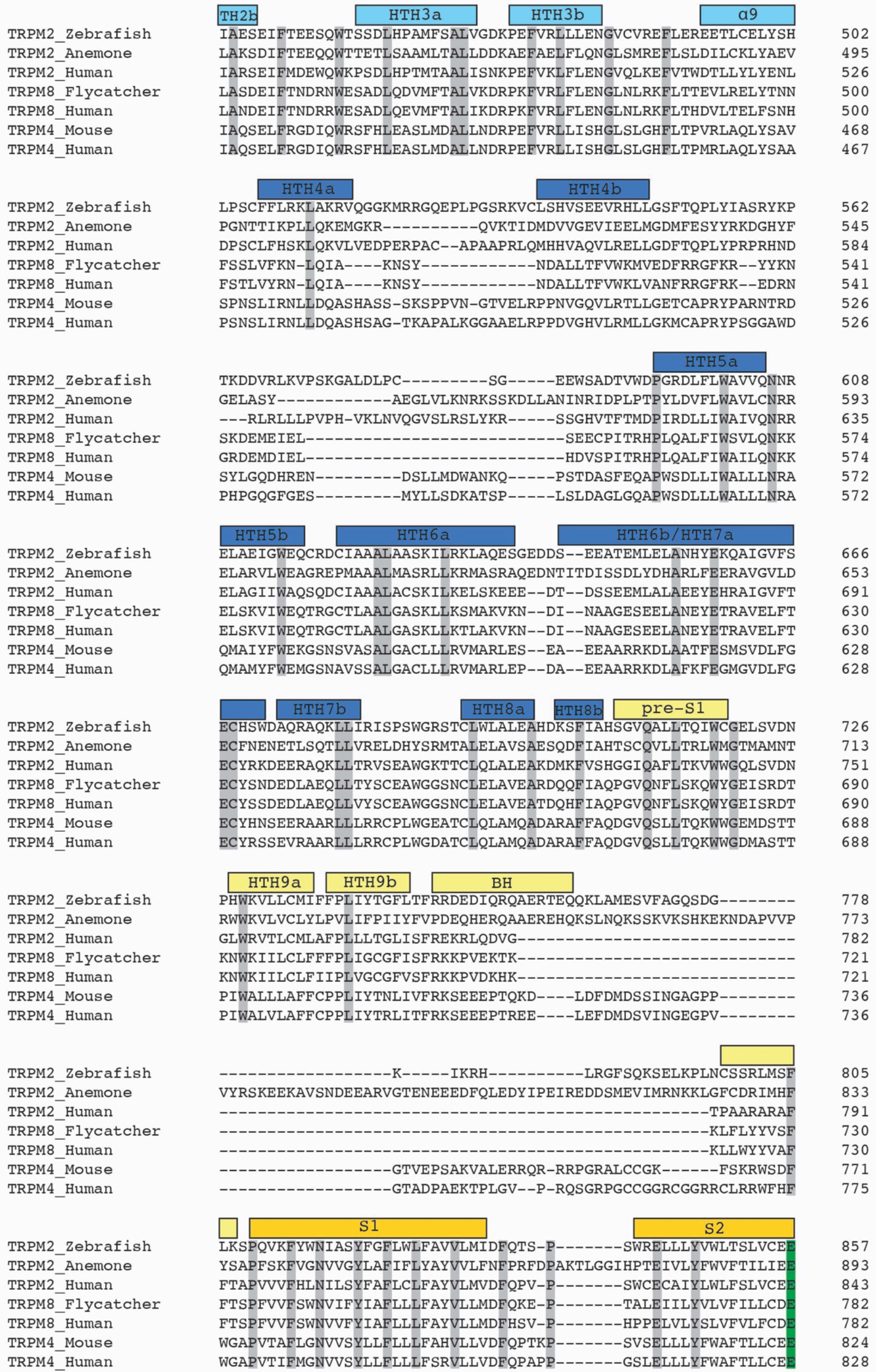

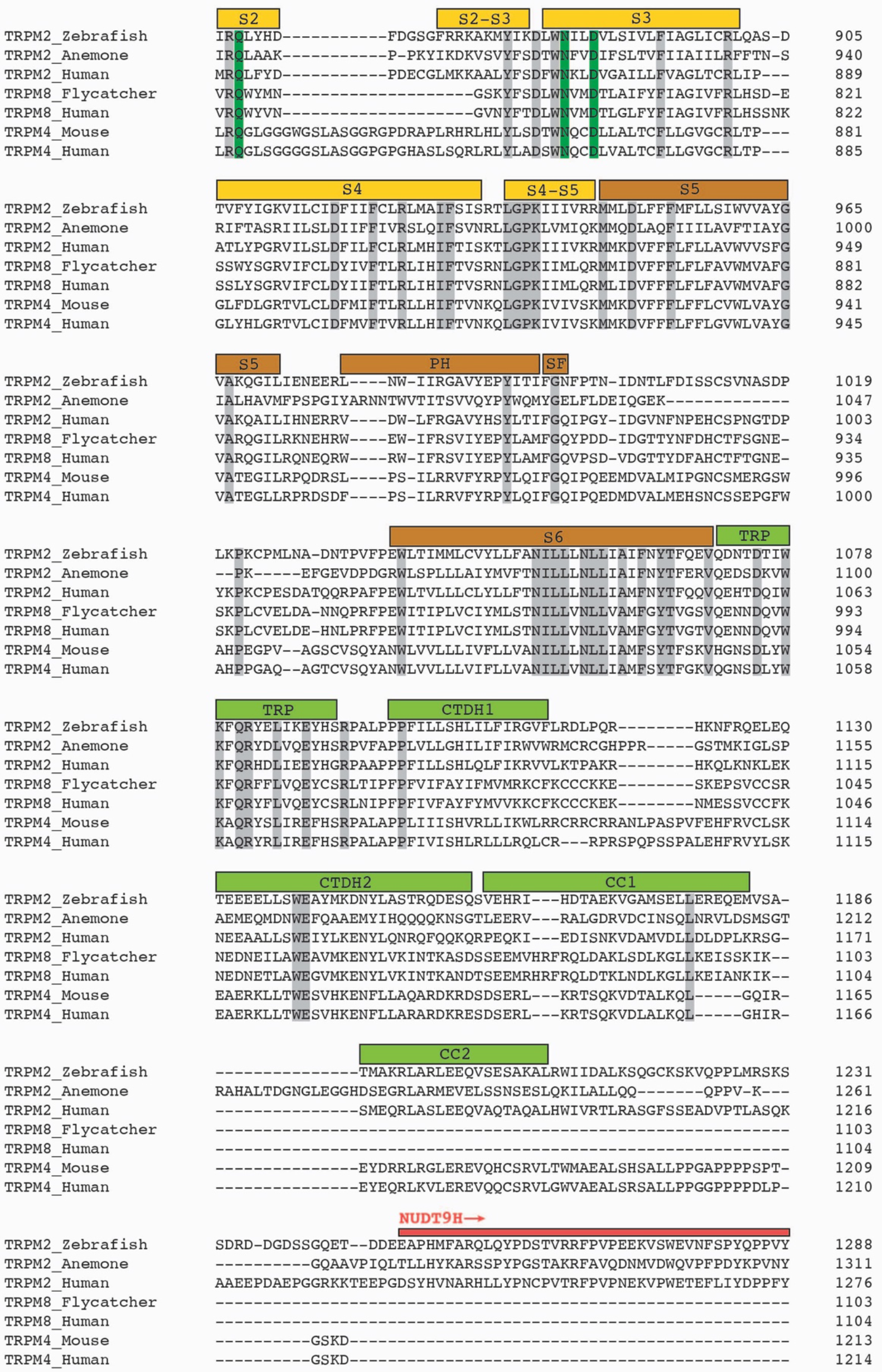

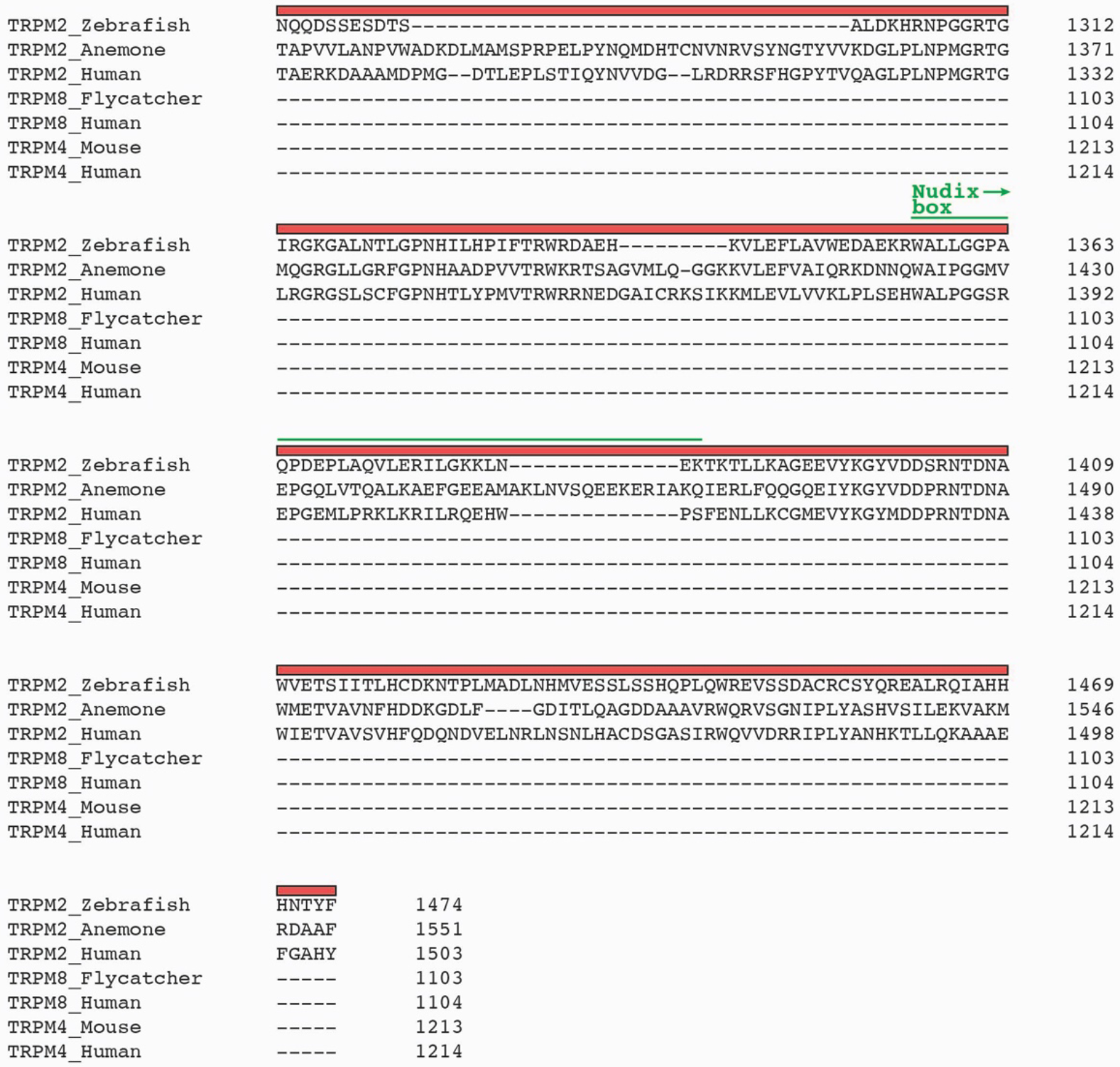

**Supplementary Figure 2.**
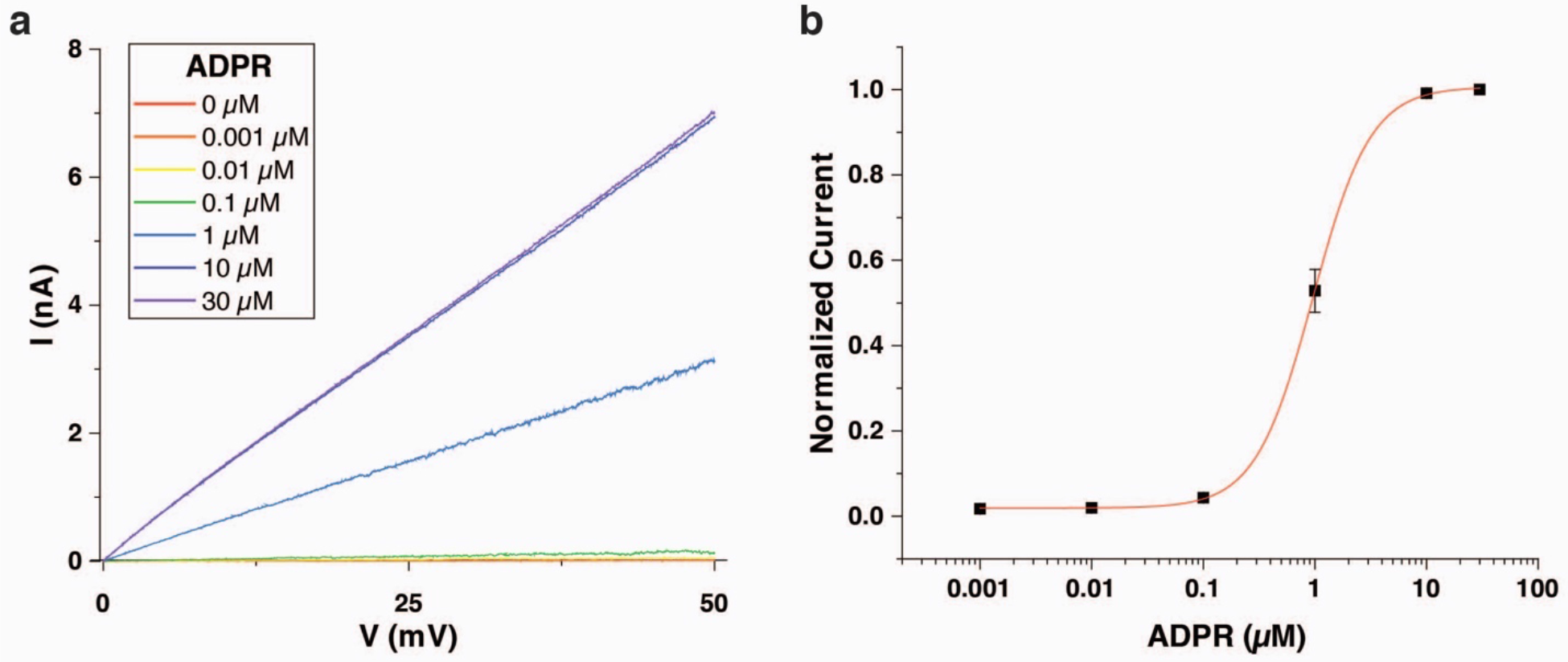
Functional characterization of the TRPM2_DR_ channel. **a**, Representative inside-out current traces recorded with repeated voltage ramps (from 0 to +50 mV; 400 ms) from HEK293T cells transfected with TRPM2_DR_ during application of 0 μM (red), 0.001 μM (orange), 0.01 μM (yellow), 0.1 μM (green), 1 μM (blue), 10 μM (indigo), and 30 μM (violet) ADPR to the inside of the patch membrane in the presence of 125 μM Ca^2+^. **b**, ADPR dose-response relationship of TRPM2_DR_ currents measured at +50 mV (V_m_ = −50 mV) generated from five samples (n = 5; biologically independent experiments) with the Hill equation. The half-maximal ADPR concentration (EC_50_) was 0.96 ± 0.01 μM with a Hill coefficient (n_H_) of 1.7 ± 0.1. Error bars represent ± SEM.

**Supplementary Figure 3.**
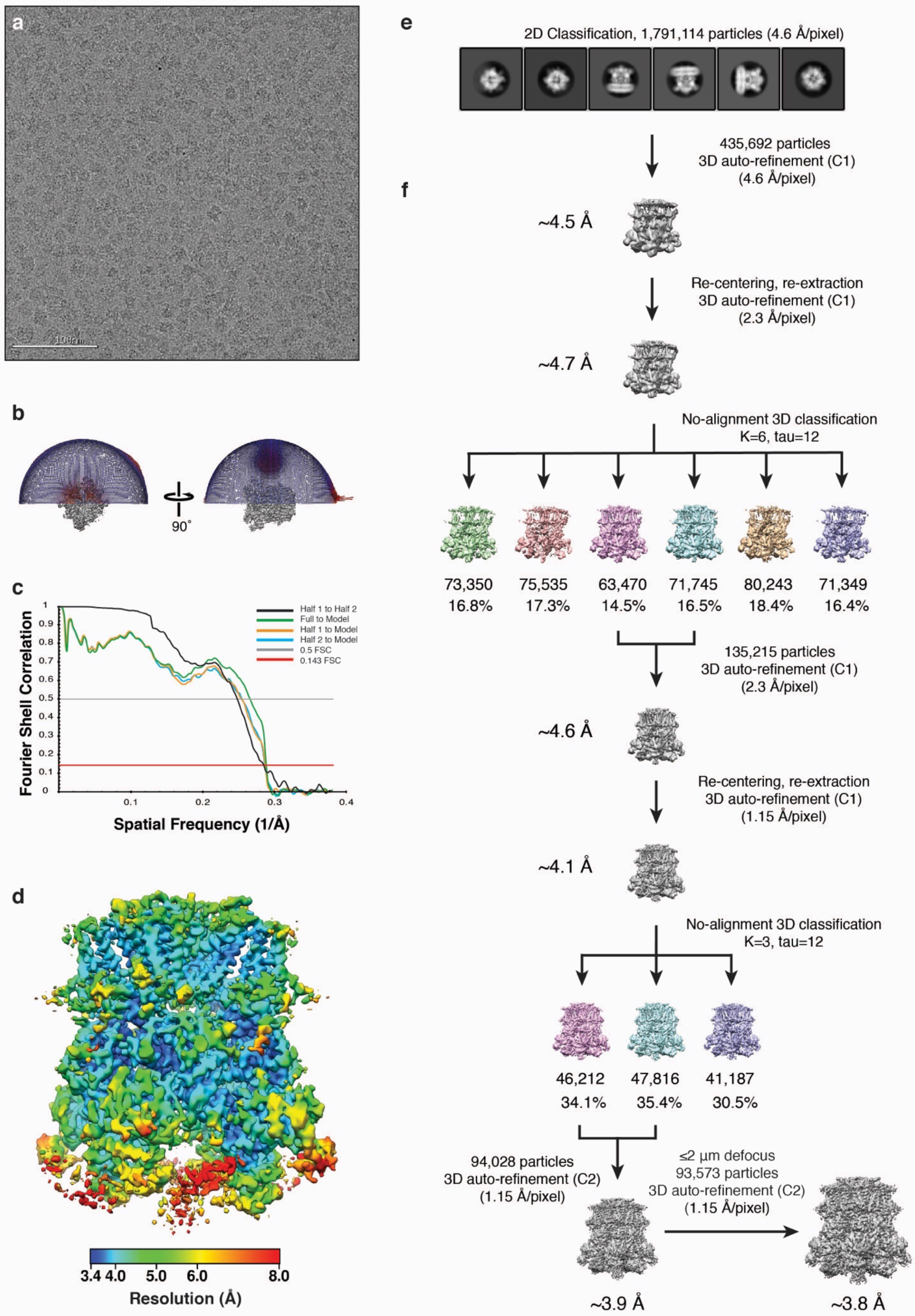
Cryo-EM data processing of TRPM2_DR_Ca2+_ structure. **a**, Representative micrograph of TRPM2_DR_Ca2+_ in vitreous ice. 3,039 movies of TRPM2 were collected over continuous carbon. **b**, Euler distribution plot for the final reconstruction. **c**, FSC curves calculated between the half maps (black line), atomic model and the final map (green line), and between the model and each half-map (orange and blue lines). **d**, Local resolution estimates of the final reconstructions calculated using BSOFT ^52^. **e**,**f**, 1,791,114 particles were extracted from aligned micrographs, Fourier binned 4 × 4, and subjected to reference-free 2D classification using RELION ^39^. Representative 2D class averages are shown (**e**). Particles comprising the “best” class averages were 3D auto-refined without symmetry to yield a ~4.5 Å resolution reconstruction. Refined particle coordinates were used for re-centering and extraction of particles. Particles were Fourier binned by 2 × 2, followed by 3D auto-refinement and no-alignment 3D classification. For each class, the number of contributing particles and percentage relative to total particles input to classification are listed below, respectively. 135,215 particles corresponding to the best-resolved classes were combined and 3D auto-refined to yield a ~4.6 Å resolution reconstruction. Refined particle coordinates were re-centered and extracted unbinned, auto-refined, and subjected to no-alignment classification to obtain a subset of 94,028 particles. Particles collected at greater than 2 μm defocus were removed to obtain a final particle stack of 93,573 particles, which was auto-refined with C2 symmetry enforced to yield a final reconstruction at ~3.8 Å resolution (**f**).

**Supplementary Figure 4.**
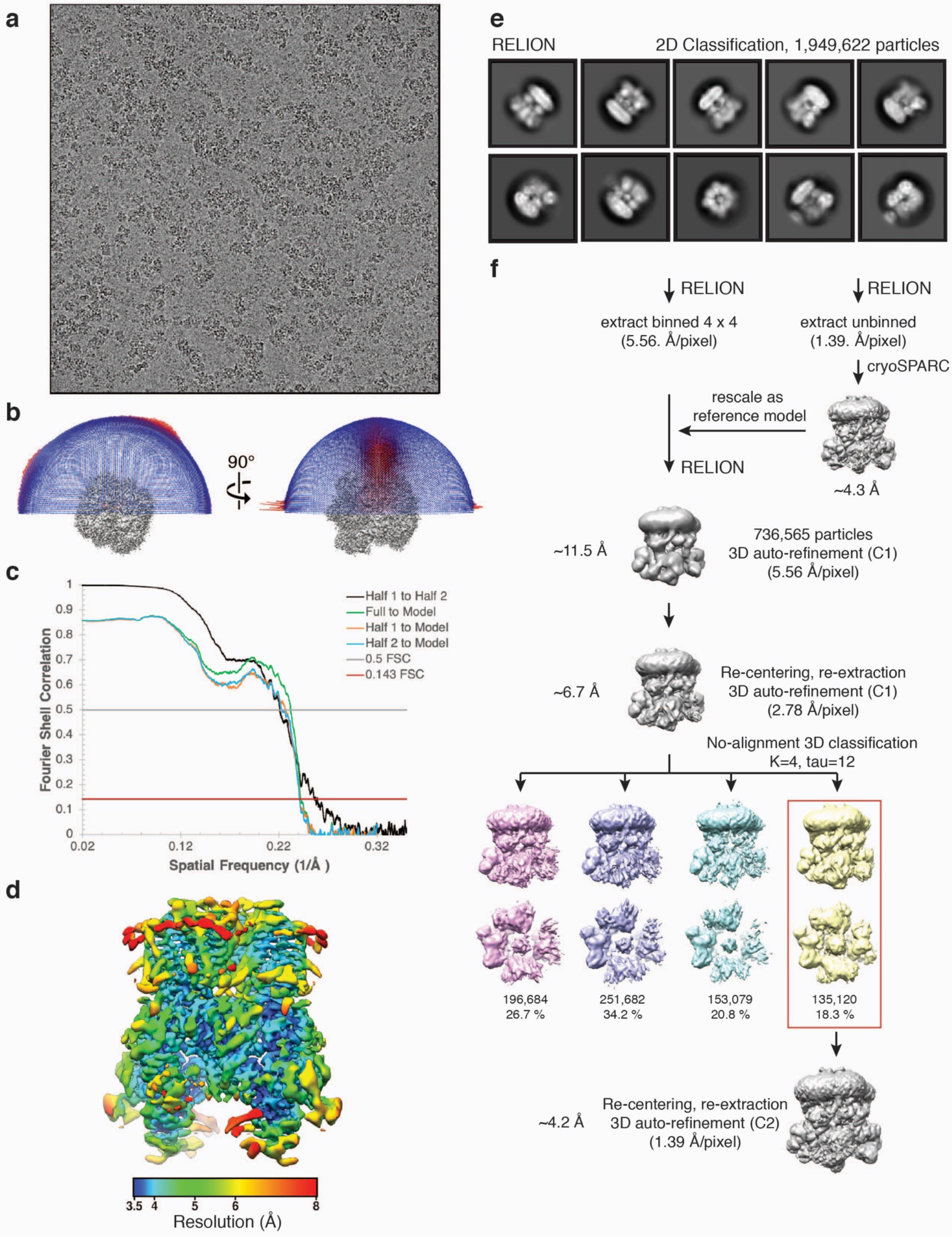
Cryo-EM data processing of TRPM2_DR_ADPR/Ca2+_ structure. **a**, Representative micrograph of TRPM2_DR_ADPR/Ca2+_ sample in vitreous ice. 2496 movies were collected. **b**, Euler distribution plot for the final reconstruction. **c**, FSC curves calculated between the half maps (black line), molecular model and the full map (green line), and between the model and each half-map (orange and blue lines). **d**, Local resolution estimation of the final reconstruction calculated using BSOFT ^52^. **e**, Particles comprising the good 2D classes were 3D auto-refined to a reconstruction of ~11.5 Å with C1 symmetry using the 3D reconstruction generated by cryoSPARC as the reference model. Refined 736,565 particles were re-extracted, re-centered, Fourier binned 2 × 2, 3D auto-refined with C1 symmetry and subjected to no-alignment 3D classification. For individual class, the number of particles and the percentage relative the total number of particles input to the classification are listed. 135,120 particles comprising the 3D class, in which the cytoplasmic domain (CD) is most well-resolved, were re-centered, re-extracted, and unbinned, and were subjected to 3D auto-refinement with C2 symmetry, yielding a final reconstruction of ~4.2 Å.

**Supplementary Figure 5.**
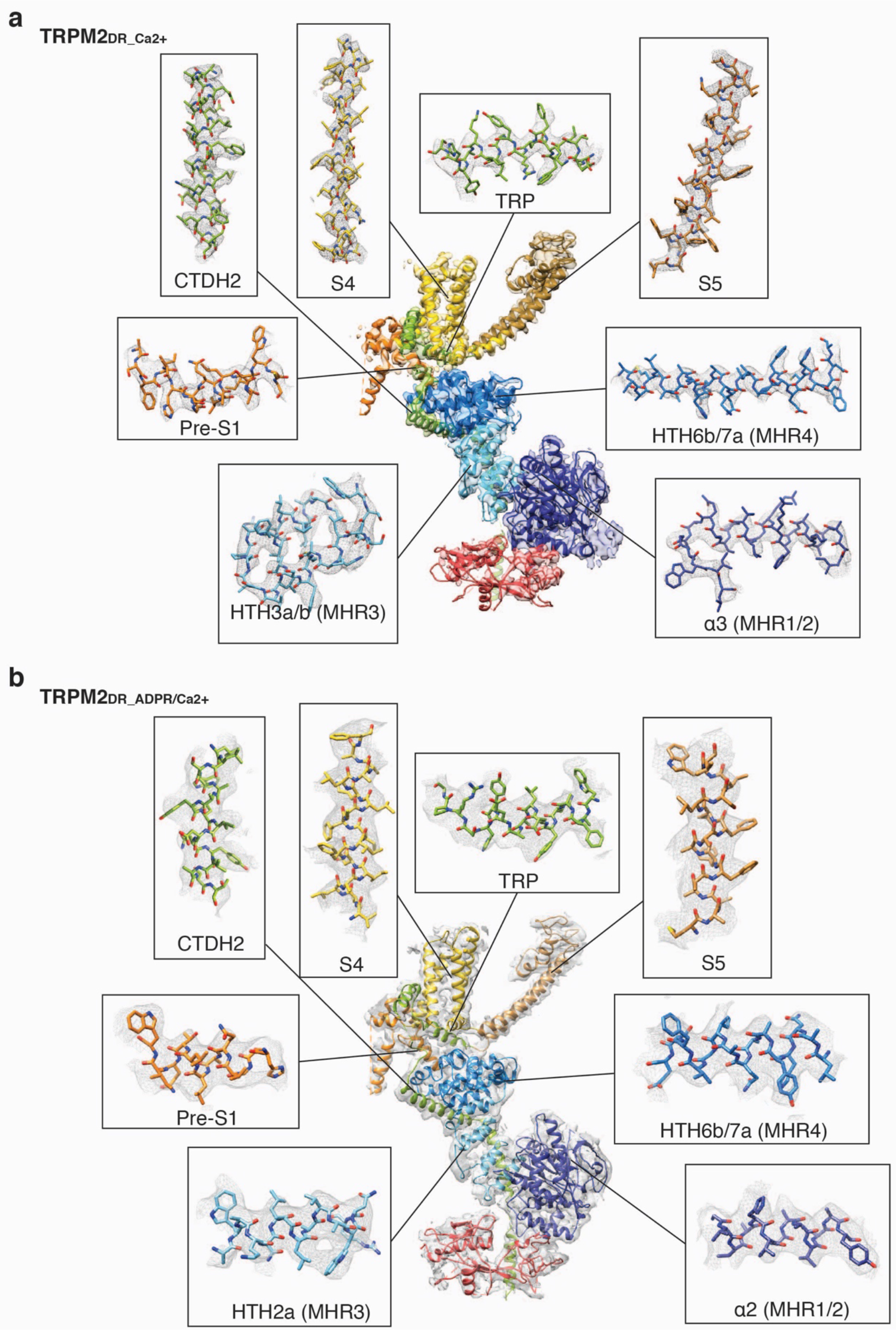
Quality of electron density of key structural elements. **a**,**b**, The structural elements in the TRPM2_DR_Ca2+_ (**a**) and TRPM2_DR_ADPR/Ca2+_ (**b**) structures are labeled and colored according to Fig. 1d and shown as sticks. The electron density is shown as a gray mesh, zoned ~2 Å around atoms in (**a**) and zoned ~2.5 Å around atoms in (**b**).

**Supplementary Figure 6.**
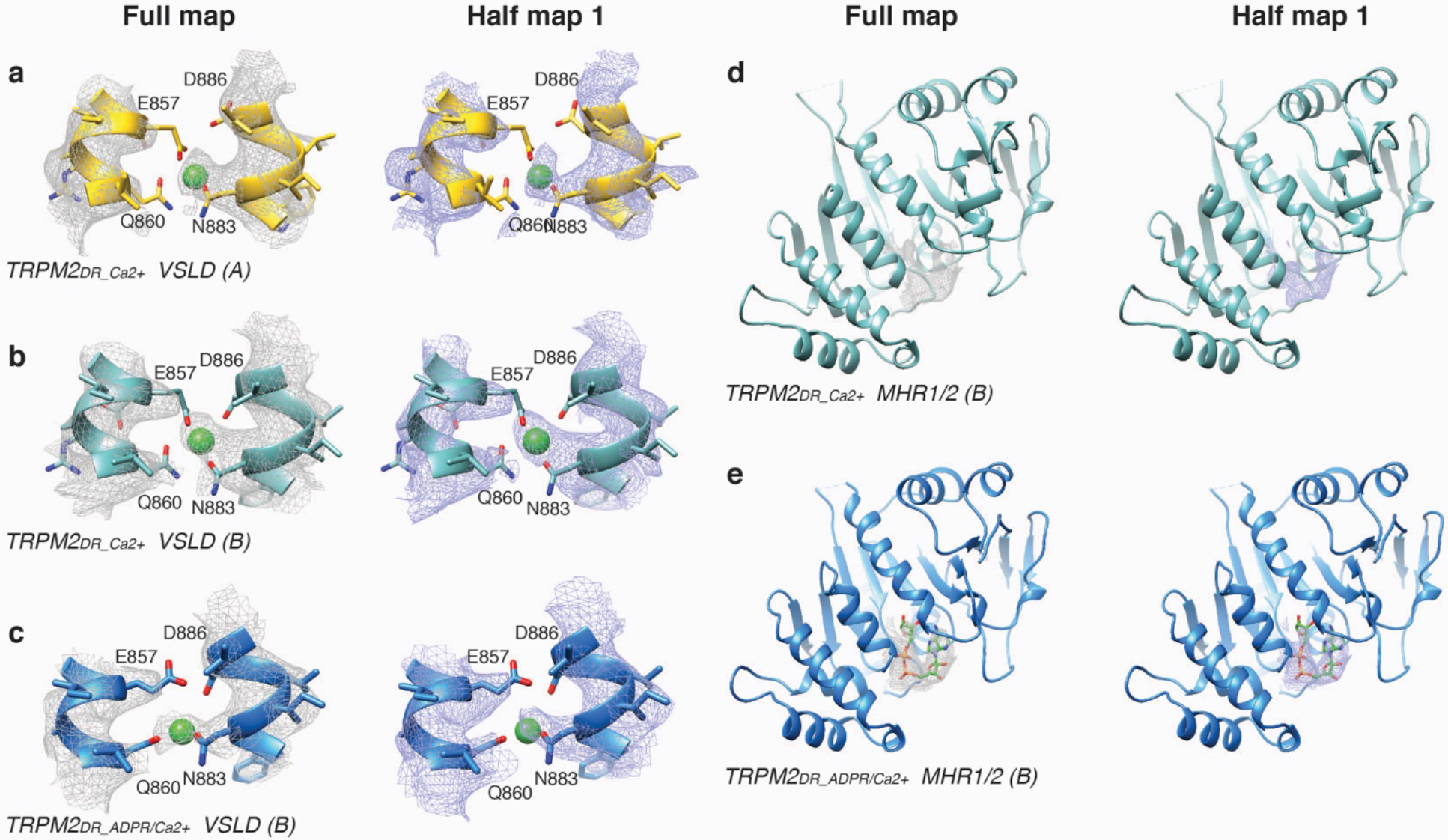
Cryo-EM densities of Ca^2+^ and ADPR in the TRPM2_DR_ structures. **a**, Cryo-EM density of the Ca^2+^ ion located in the VSLD cavity of the protomer A in the TRPM2_DR_Ca2+_ structure, in the full map (left, grey mesh) and half map 1 (right, purple mesh) zoned ~2.5 Å around atoms at 0.045 and 0.025 thresholding. **b**, Cryo-EM density of the Ca^2+^ ion located in the VSLD cavity of the protomer B in the TRPM2_DR_Ca2+_ structure, in the full map (left, grey mesh) and half map 1 (right, purple mesh) zoned ~2.5 Å around atoms at 0.045 and 0.025 thresholding. **c**, Cryo-EM density of the Ca^2+^ ion located in the VSLD cavity of the protomer B in the TRPM2_DR_ADPR/Ca2+_ structure, in the full map (left, grey mesh) and half map 1 (right, purple mesh) zoned ~2.5 Å around atoms at 0.05 thresholding. **d,** Putative EM density resembling ADPR located in the MHR1/2 domain of the protomer B in the TRPM2_DR_Ca2+_ structure, in the full map (left, grey mesh) and half map 1 (right, purple mesh) zoned ~2.5 Å around atoms at 0.05 and 0.035 thresholding. **e**, Cryo-EM density of the ADPR located in the MHR1/2 domain of the protomer B in the TRPM2_DR_ADPR/Ca2+_ structure, in the full map (left, grey mesh) and half map 1 (right, purple mesh) zoned ~2.5 Å around atoms at 0.045 thresholding.

**Supplementary Figure 7.**
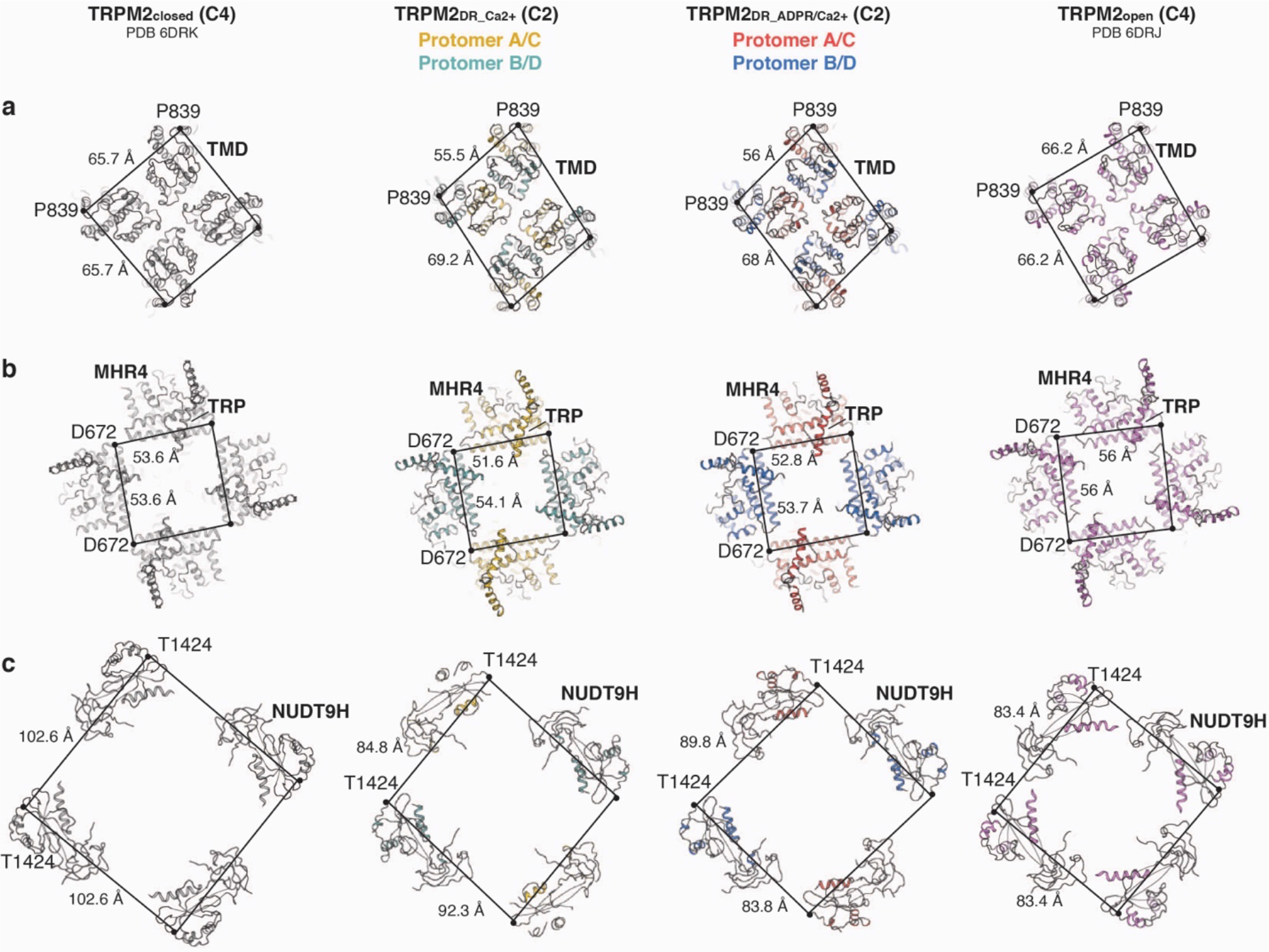
Comparison of the channel symmetry in TRPM2_DR_ structures. **a-c**, Extracellular views of the TMDs (**a**), sliced between the TMDs and the CDs (**b**), and between the middle and bottom layer of the CDs (**c**) in the TRPM2_closed_, TRPM2_DR_Ca2+_, TRPM2_DR_ADPR/Ca2+_, and TRPM2_open_ structures, comparing the two-fold symmetry in the TRPM2_DR_Ca2+_ and TRPM2_DR_ADPR/Ca2+_ structures with the four-fold symmetry in the TRPM2_closed_ and TRPM2_open_ structures.

**Supplementary Figure 8.**
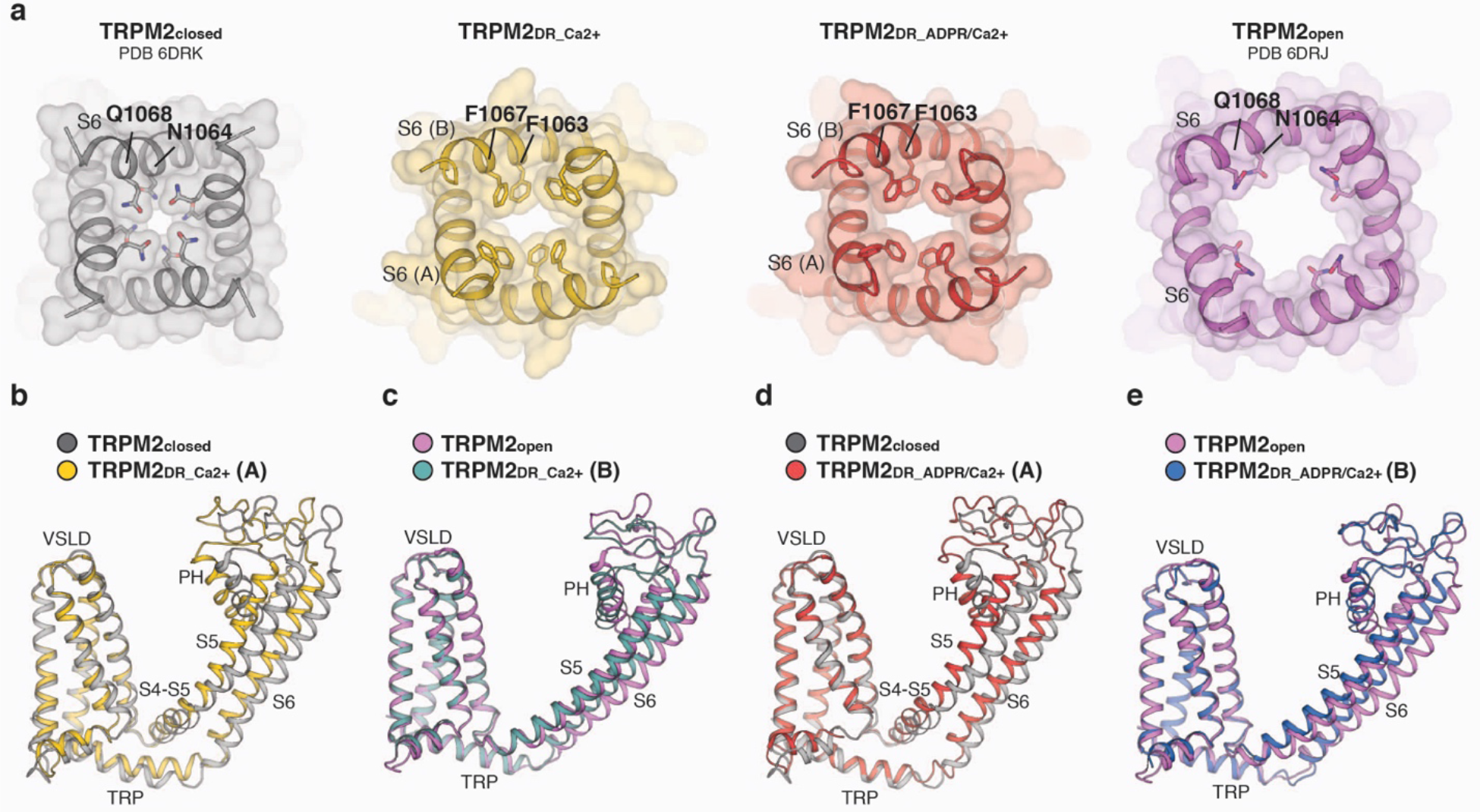
Comparison of the transmembrane channel domain (TMD) in TRPM2_DR_ structures. **a**, Cartoon and surface representations of the S6 gate viewed from the intracellular side. In the TRPM2_closed_ (grey) and TRPM2_open_ (violet) structures, residues N1064 and Q1068 form the narrowest restriction points, while residues F1063 and F1067 on S6 form the intracellular gate in the TRPM2_DR_Ca2+_ (yellow) and TRPM2_DR_ADPR/Ca2+_ (red) structures. **b**, the TMD of protomer A in the TRPM2_DR_Ca2+_ structure (yellow) resembles that of the TRPM2_closed_ structure (silver). **c**, the TMD of protomer B in the TRPM2_DR_Ca2+_ structure (teal) resembles that of the TRPM2_open_ structure (violet). **d**, the TMD of protomer A in the TRPM2_DR_ADPR/Ca2+_ structure (red) resembles that of the TRPM2_closed_ structure (silver). **e**, the TMD of protomer B in the TRPM2_DR_ADPR/Ca2+_ structure (blue) resembles that of the TRPM2_open_ structure (violet).

**Supplementary Figure 9.**
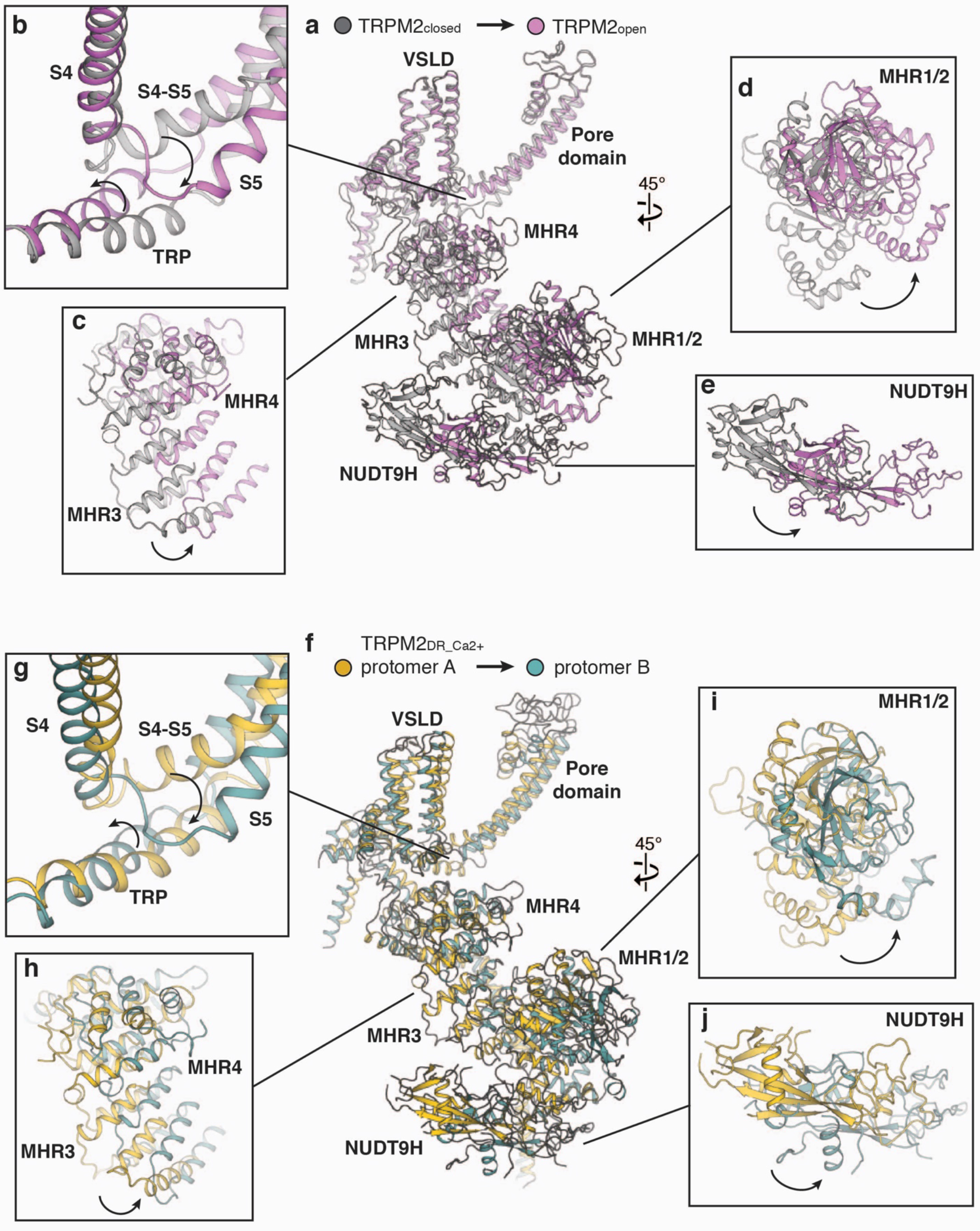
Conformational rearrangements during TRPM2 channel gating. **a**, Viewed from the membrane plane, cartoon representations of TRPM2_closed_ and TRPM2_open_ protomers aligned at the TMDs, showing how conformational changes in the CD could be propagated to TMD during channel activation. Protomers of TRPM2_closed_ and TRPM2_open_ are colored in silver and violet respectively. **b-e**, Close-up views of the structural rearrangements of TRP domain and S4-S5 linker (**b**), MHR3-4 (**c**), MHR1/2 (**d**), and the NUDT9H domain (**e**) depicted in (**a**). **f**, Viewed from the membrane plane, cartoon representations of protomers A and B of TRPM2_DR_Ca2+_ structure aligned at the TMDs, showing conformational changes propagated from CD to TMD resemble those depicted in (**a**). Protomers A and B are colored in gold and teal respectively. **g-j**, Close-up views of the structural rearrangements of TRP domain and S4-S5 linker (**g**), MHR3-4 (**h**), MHR1/2 (**i**), and the NUDT9H domain (**j**) depicted in (**f**).

**Supplementary Figure 10.**
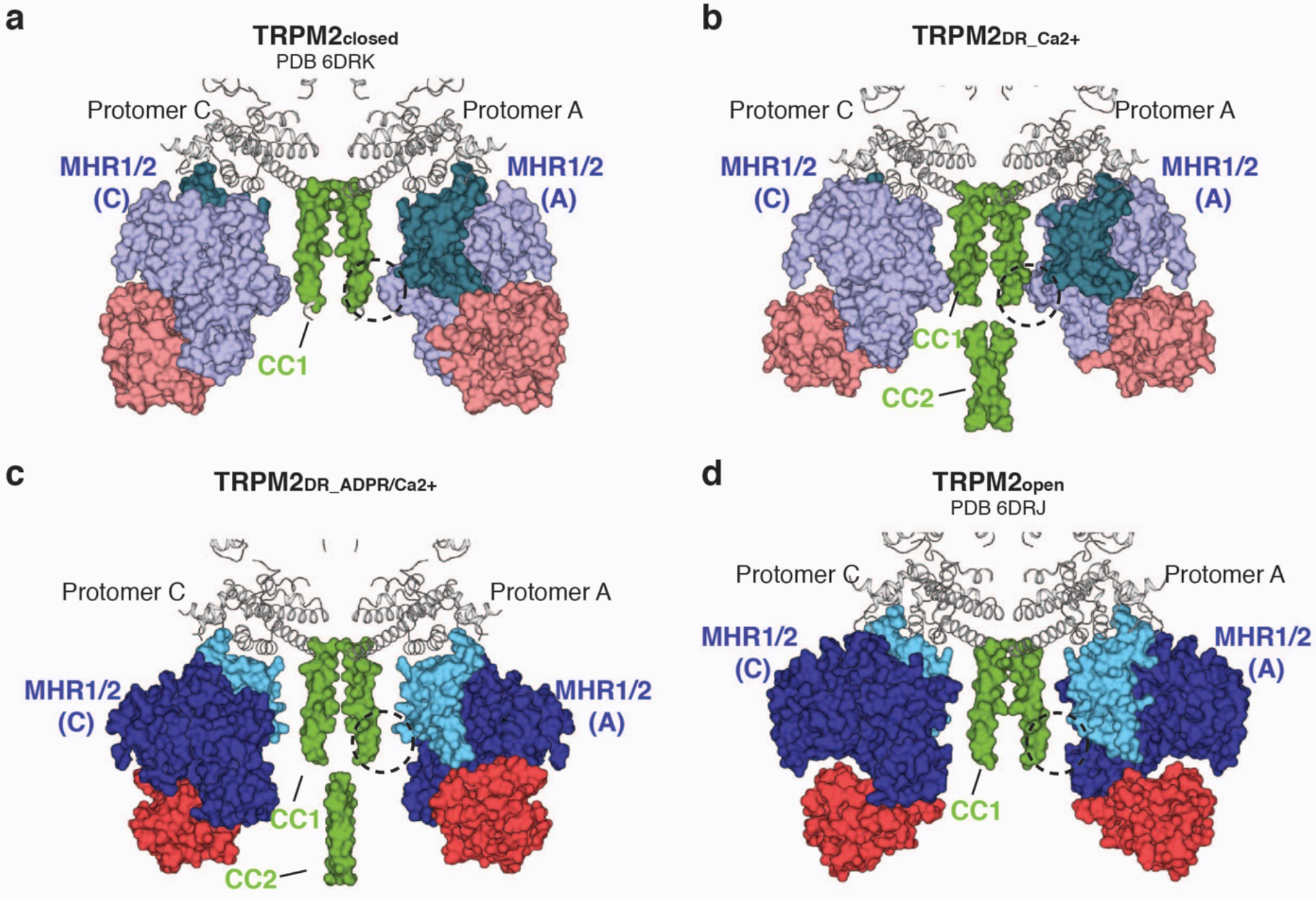
Comparison of the interfacial contact between CC1 and MHR1/2 domain in different TRPM2_DR_ structures. **a-d**, Cross-section views of opposing protomers A and C in the published TRPM2_closed_ structure (**a**, PDB 6DRK), in TRPM2_DR_Ca2+_ (**b**) and TRPM2_DR_ADPR/Ca2+_ (**c**) structures from the current study, and in the published TRPM2_open_ structure (**d**, PDB 6DRJ), showing the detachment of MHR1/2 domain from CC1 in the TRPM2_DR_ADPR/Ca2+_ (**c**) and TRPM2_open_ (**d**) structures, in contrast with the association between the two domains in the TRPM2_closed_ (**a**) and TRPM2_DR_Ca2+_. (**b**) structures.

